# Spread of a single superclone drives insecticide resistance in *Acyrthosiphon kondoi* across an invasive range

**DOI:** 10.1101/2024.12.16.628636

**Authors:** Joshua A. Thia, Benjamin J. Hunt, Shuchao Wang, Bartlomiej J Troczka, Evatt Chirgwin, Courtney J. Brown, Rumi Sakamoto, Monica Stelmach, Kelly Richardson, Leonhard S. Arinanto, Ashritha P. S. Dorai, Chinmayee Joglekar, Qiong Yang, Marielle Babineau, Chris Bass, Paul A. Umina, Ary A. Hoffmann

**Affiliations:** School of BioSciences, The University of Melbourne, Melbourne, Victoria, Australia; Centre for Ecology and Conservation, University of Exeter, Penryn, Cornwall, UK; Cesar Australia, Brunswick, Victoria, Australia

## Abstract

Populations under similar selection pressures may adapt via parallel evolution or dispersal of advantageous alleles. Here, we investigated insecticide resistance in the invasive blue-green aphid, *Acyrthosiphon kondoi*, which reproduces clonally in Australia and has rapidly developed resistance across geographic locations. Using genomic, transcriptomic, and experimental approaches, we explored the evolutionary origins and molecular mechanisms of resistance. We developed the first nuclear genome assembly for *A. kondoi* (443.8 Mb, 28,405 annotated genes, BUSCO score 97.5%) and a partial mitochondrial assembly (11,598 bp). All resistant strains shared a common ancestor, supporting the spread of a resistant ‘superclone’ lineage that is distinct from susceptible strains. Resistance was associated with over-expression of an esterase gene that was homologous to E4/FE4 esterases in other aphid pests that are linked to resistance.

Functional experiments in *Drosophila melanogaster* confirmed a causal role of this E4- like esterase in resistance to organophosphates, carbamates, and pyrethroids. These findings highlight how clonal dispersal and insecticide overuse can transform local adaptation into a widespread pest management issue. Our results suggest a parallel macroevolutionary response to insecticide selection in *A. kondoi* and other aphid species at the gene family level, but with a distinct regulatory mechanism in *A. kondoi*. Given the rapid spread of the resistant superclone, alternative management strategies, including expanded chemical control options and enhanced biological control, are urgently needed to mitigate this growing pest problem.

## Introduction

Understanding variation in the rise and spread of adaptive traits and their underlying genes is a key issue in evolutionary biology (Cohan and Hoffmann 1989; Korona et al. 1994; Colosimo et al. 2005). Populations or species exposed to similar selective pressures may evolve through parallel changes (independent evolutionary events). Parallelism might occur only at the phenotypic level (Roda et al. 2013; Magalhaes et al. 2020; Yang et al. 2020; Sprengelmeyer and Pool 2021), or it may extend to the genotypic level, where the same allelic changes are involved. The latter is more likely if populations share standing genetic variation or if there are genetic constraints on evolving adaptive phenotypes (Colosimo et al. 2005; Yeaman et al. 2018; Magalhaes et al. 2020; Montejo- Kovacevich et al. 2022). Gene flow can also spread adaptive alleles across areas with similar selective pressures (Endersby-Harshman et al. 2020; Tepolt et al. 2021; Zhang et al. 2023). Indeed, gene flow can assist adaptation to new environmental stressors by introducing adaptive variation into local populations (Chevillon et al. 1999; Bontrager and Angert 2019). Disentangling these processes sheds light on the repeatability of evolution and the origins of adaptive variation in local populations.

Clonal organisms are often considered to have lower adaptive capacity because they lack recombination to produce adaptive allelic combinations (Muller 1932; Barraclough et al. 2003). However, when clonal diversity arises from mutation accumulation or rare sexual reproduction, it is possible for selection to drive clonal lineages with adaptive alleles to high frequency, often at the expense of other lineages (Muller 1932; Gerrish and Lenski 1998; Barraclough et al. 2003). Clonal organisms therefore provide a unique context for tracking the origin and trajectory of adaptive traits and alleles, as these are directly linked to specific (clonal) genetic backgrounds. For example, in the cystic fibrosis-associated bacterium *Pseudomonas aeruginosa*, recurrent selective sweeps have been associated with parallel genetic, transcriptional, and phenotypic changes for adaptive pathogenicity among clonal lineages (Huse et al. 2010; Feliziani et al. 2014; Caballero et al. 2015). In the parasitic protozoan, *Toxoplasma gondii*, dominance of three clonal lineages in North America and Europe has been attributed to strong selection on key virulence alleles associated with each of these lineages (Howe and Sibley 1995; Khan et al. 2009). Studies on *Daphnia* species have also shown clear, predictable patterns of changes in clone frequencies due to variation in competitive abilities of clonal lineages under different environmental conditions (Korpelainen 1986; Steiner et al. 2016; Lyberger and Schoener 2023). And in yeast, tracking of clonal lineages has illustrated how the accumulation of beneficial mutations within clones leads to the dominance of highly fit genotypes (Nguyen Ba et al. 2019).

Insecticide usage in agriculture provides an interesting context for studying adaptation to similar selective environments. Pest species are often exposed to the same insecticide across different populations as part of chemical control strategies (Walsh et al. 2018; Endersby-Harshman et al. 2020; Singh et al. 2021; Thia, Korhonen, et al. 2023). Because insecticides exert their toxicity through very specific protein targets and molecular interactions, there is strong precedence for joint phenotypic and genotypic parallelism in insecticide resistance traits. This has been demonstrated repeatedly, leading to parallel and convergent evolution of some mutations in these target proteins that prevent insecticide inhibition – so called ‘target-site mutations’ leading to amino acid substitutions (Fournier 2005; Davies et al. 2007). Such parallelism can occur both at the microevolutionary scale among populations (Walsh et al. 2018; Thia, Korhonen, et al. 2023), and at the macroevolutionary scale, leading to convergence among species (Feyereisen et al. 2015). However, alternate mechanisms might also evolve, such as shifts in the expression or efficacy of detoxification proteins, and polygenic resistance can lead to non-parallel genetic mechanisms of insecticide resistance among populations (Field and Devonshire 1998; Crossley et al. 2017; Yang et al. 2020; Thia, Korhonen, et al. 2023).

Although clonal reproduction is uncommon in animal populations, it tends to occur at a relatively higher frequency among invertebrates in agricultural environments (Hoffmann et al. 2008). Aphids (Order: Hemiptera) are a major agricultural arthropod pest: they are sap-sucking insects that damage plants through direct feeding and virus transmission (Blackman and Eastop 2017). Aphid populations are often composed of holocyclic (sexual or asexual, host-alternating) clones, particularly in temperate regions with cold winter temperatures, but they often become anholocyclic (asexual, nonhost-alternating) in more benign climates (Rispe et al. 1998; Rispe and Pierre 1998; Williams and Dixon 2017). Clonal reproduction can dramatically shape the evolutionary trajectory of aphid populations by favouring the dominance of ‘superclones’. These are clonal lineages that have acquired highly advantageous traits (through genetic mutation or endosymbiont acquisition) that allow them to spread across broad geographic extents (Gilabert et al. 2015; Godefroid et al. 2024; Mahieu et al. 2024). For agriculturally important aphid species, insecticide selection can facilitate the spread of superclones carrying insecticide resistance mutations. For example, in the peach-potato aphid, *Myzus persicae*, although different genetic lineages carrying resistance have evolved around the world (Singh et al. 2021), local sweeps of specific superclones have occurred in places like Australia (Umina et al. 2014; de Little et al. 2017; Ward et al. 2024). Similarly, in the wheat aphid, *Sitobion avenae*, a single insecticide resistant clonal lineage has become dominant in the UK, reducing local clonal diversity (Morales-Hojas et al. 2020).

In this study, we investigated the recently evolved case of insecticide resistance in the blue green aphid, *Acyrthosiphon kondoi*, in its invasive Australian range. This aphid is native to Asia but has become an agricultural pest across the world, including the USA, South America, Africa, and Australia (Ryalls et al. 2013). It is a pest of alfalfa and leguminous grains crops such as lentils and lupins (Humphries et al. 2012; Ryalls et al. 2013). In Australia, and in other parts of its invasive range, *A. kondoi* reproduction is mostly asexual (Ryalls et al. 2013), with sexual reproduction often occurring in its native range, though data are limited (Kawada 1992). Historically, there are few reports of insecticide resistance in *A. kondoi*. One of the only cases is described in unpublished work from the USA (Natwick et al. 2014). Since then, there have been no documented cases globally until the recent discovery of organophosphate, carbamate, and pyrethroid resistance in Australian *A. kondoi* populations in 2020 (Chirgwin et al. 2023). This has led to control failures and subsequent economic damage to Australian alfalfa growers (Chirgwin et al. 2023). Two outstanding issues are whether the resistance has a single evolutionary origin in Australian *A. kondoi*, and what genes contribute to this adaptive trait.

Here, we consider the evolved resistance in Australian *A. kondoi* to study the evolutionary origins of adaptation and its genetic basis. We began with a genomics approach to address two initial objectives: (1) to determine whether resistant strains are of the same or different clonal background; and (2) to identify putative genes that might underpin resistance. Our finding of a candidate detoxification gene, an E4-like esterase, led us to address two additional objectives: (3) to validate the differential expression patterns of the candidate E4-like esterase among resistant and susceptible strains; and (4) to validate the potential insecticide detoxification function provided by the candidate E4-like esterase using transgenic *Drosophila*.

## Results

### Aphid strains

Our study leveraged a suite of field collections of clonal strains between 2020 and 2024, as well as a laboratory clonal strain that had been isolated in 1999 before resistance was reported in the field. The clonal strains came from a variety of plant hosts and were broadly distributed across the southern regions of Australia (Table 1 and Figure 1a). We first developed a set of eight core clonal isofemale strains using artificial selection with the organophosphate insecticide, chlorpyrifos. These clonal isofemale strains formed the core of our main experiments. Susceptible isofemale strains included Kellerberrin, Eudunda, Jung and Moulamein, and resistant isofemale strains included Keith, Temora, Bangham and Coombe. Success of selection was validated with follow-up screens using chlorpyrifos (Supplementary Table S1). Previous work has demonstrated that resistant strains of *A. kondoi* exhibit general resistance to carbamates, organophosphates and pyrethroids (Chirgwin et al. 2023), so we did not screen our core strains against other insecticides. We later broadened our geographic sampling by sampling single individuals from field strains. The phenotypes for these field strains had been assessed in a separate study (Evatt Chirgwin, unpublished data), but we did not specifically impose the same artificial selection treatment on them as we had with our core strains.

**Figure 1.**
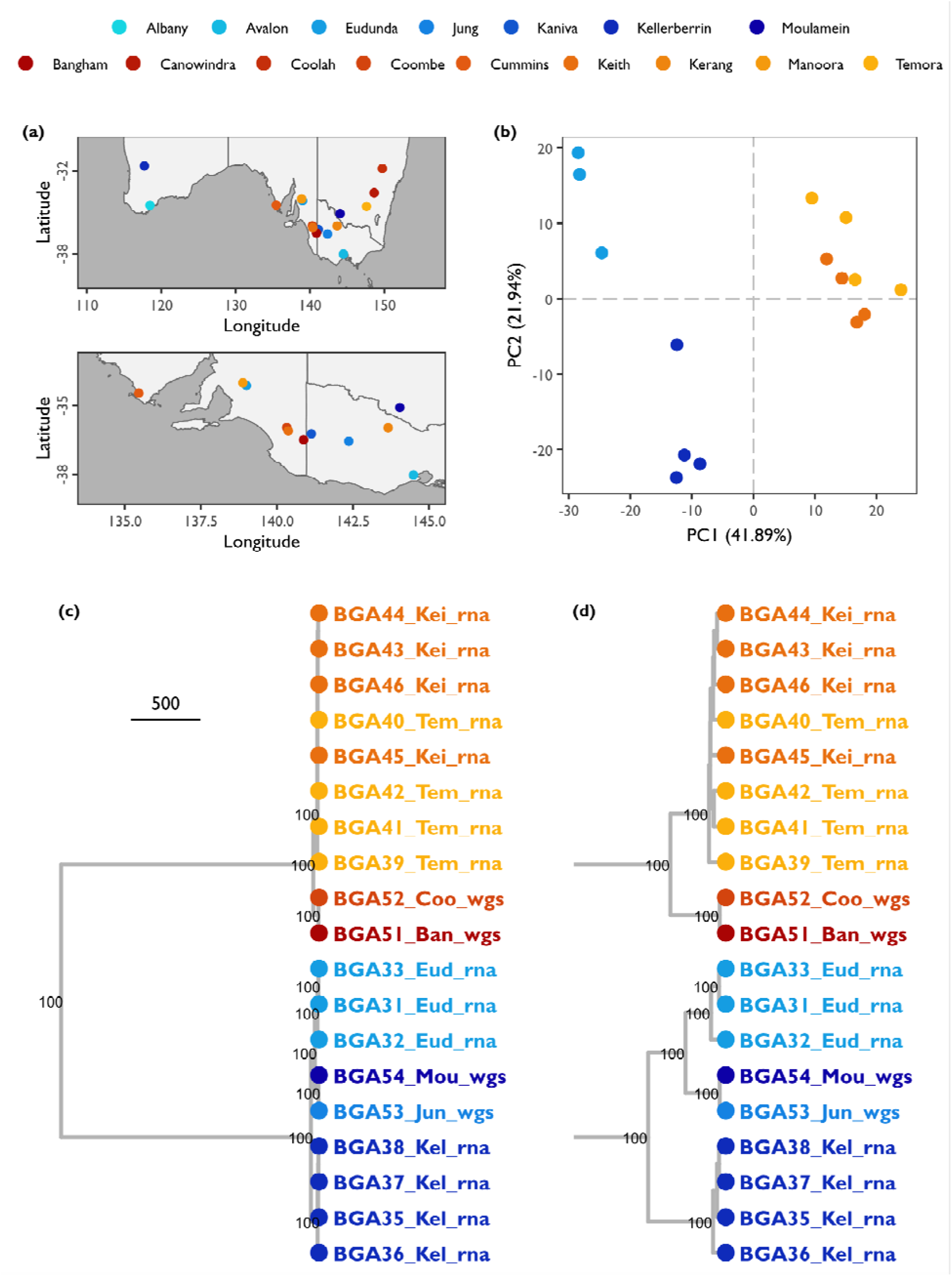
Transcriptomic and genetic comparisons of *Acyrthosiphon kondoi* strains examined in this study. The susceptible strains include Eudunda (Eud), Kellerberrin (Kel), Jung (Jun) and Moulamein (Mou). The resistance strains include Bangham (Ban), Coombe (Coo), Keith (Kei) and Temora (Tem). (a) Source locations of each strain in Australia. The top panel provides a broader geographic view of eastern and western regions, whereas the bottom panel provides a finer geographic view of southern regions. (b) Transcriptomic profiles of the Eudunda, Kellerberrin, Keith and Temora strains projected into principal component space. Percentages in parentheses on the x-axis and y-axis capture the amount of variance explained by the first and second PC axis, respectively. (c) A UPGMA tree constructed from allelic distances among samples of each strain. Branch nodes are labelled with bootstrap values >80%. Samples follow the naming convention ‘ID_Strain_Method’, where ‘ID’ is a unique sample identifier, ‘Strain’ is one of the eight sampled strains, and ‘Method’ is one of ‘rna’ for transcriptome sequencing or ‘wgs’ for whole-genome sequencing. (a–c) Points are coloured by strain, see the legend.

**Table 1.**
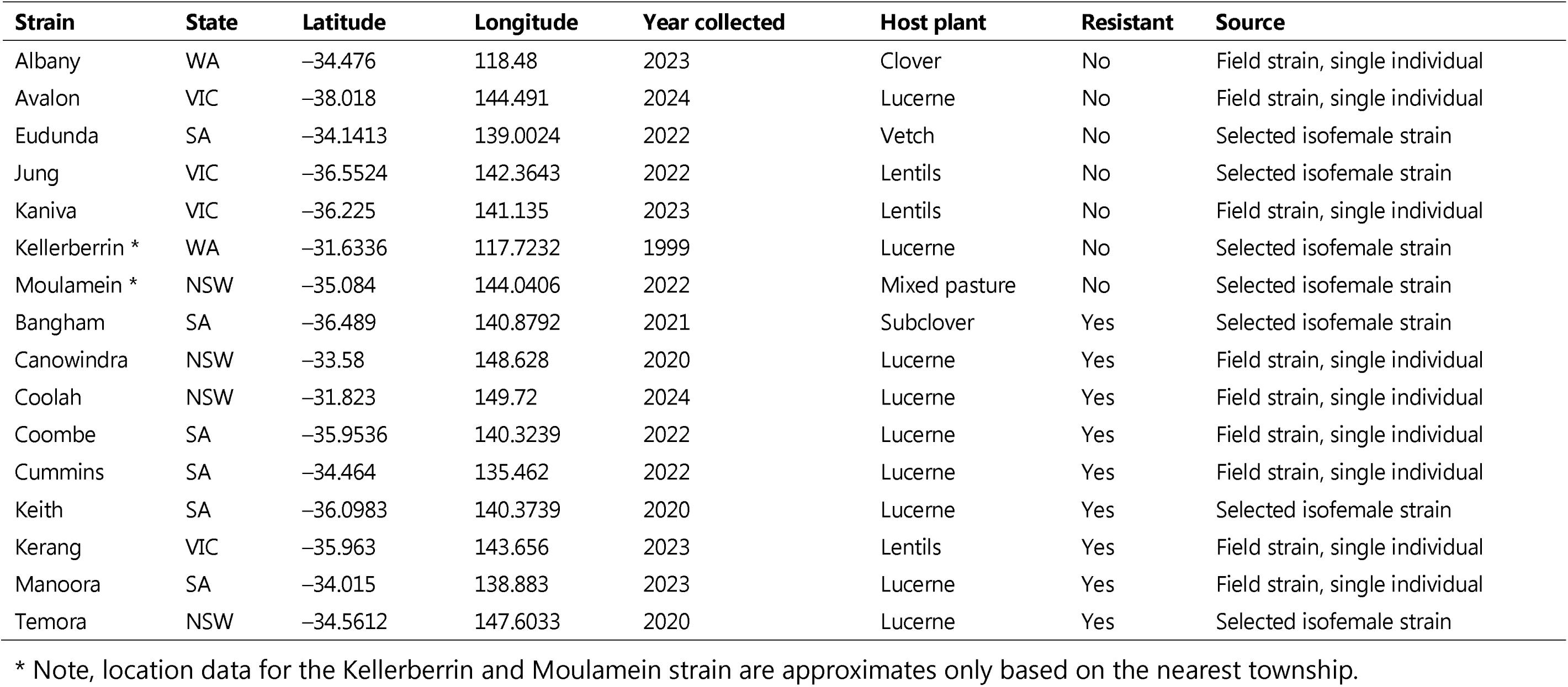
Clonal *Acyrthosiphon kondoi* strains examined in this study.

### Genome assembly and annotation

To investigate the evolution of insecticide resistance in *A. kondoi*, we generated the first nuclear reference genome assembly (NCBI Genome Accession JAUMJG000000000) and partial mitochondrial reference genome assembly (NCBI Nucleotide Accession PV793158) for this species using the Kellerberrin strain. We assembled these genomic references with a combination of whole-genome Illumina short-reads and whole-genome Nanopore long-reads, and annotated them using transcriptome Illumina short-reads.

#### Nuclear genome

Our final nuclear genome assembly was 443.8 Mb, comprising 615 contigs and 37 scaffolds, an L50 of 70 sequences, an N50 of 1.8 Mb, an L90 of 300 sequences, and an N90 of 0.3 Mb. The mean assembled sequence length was 0.7 Mb, with a minimum of 0.006 Mb and a maximum of 9.7 Mb. Repetitive content comprised 30% of the assembly. We annotated 28,198 genes. The completeness of our assembly was high based on BUSCO scores calculated for arthropod single-copy orthologues (genome/protein sequences): 97.9%/97.5% of arthropod single-copy orthologous genes were present, with 91.3%/91.7% present as single copies and 6.6%/5.8% present as duplicate copies. There were 0.7%/0.9% fragmented genes and 1.4%/1.6% missing genes. A self-to-self alignment produced a generally clean dotplot, indicating that our assembly had limited redundancy (Supplementary Figure S1a).

Our *A. kondoi* nuclear genome assembly has similar characteristics to genomes developed for the congeneric pea aphid, *A. pisum*, available on GenBank, which is the most closely related species with a chromosome-level genome assembly. The *A. pisum* GCF_000142985.2 scaffold-level assembly is 541.7 Mb, comprising 60,595 contigs and 23,924 scaffolds, with 20,940 annotated genes. The *A. pisum* GCF_005508785.2 chromosome-level assembly is 533.6 Mb, comprising 66,716 contigs and 21,227 scaffolds arranged into 4 chromosomes, with 20,307 annotated genes. We used alignments of our *A. kondoi* contigs against the *A. pisum* chromosome-level assembly (GCF_005508785.2) to generate pseudo-chromosomes for *A. kondoi*, which were then remapped these back to the *A. pisum* chromosomes. This analysis suggested that there may be considerable overlap in homologous sequences between our *A. kondoi* assembly and the *A. pisum* chromosome-level assembly (Supplementary Figure S1b). There may also be a reasonable amount of synteny between these congeneric aphids (Supplementary Figure S2). However, additional data (e.g., Hi-C sequencing) would provide greater power to test synteny.

#### Mitochondrial genome

Our mitogenome reference was a partial sequence comprising 11,598 bp. This reference contained most of the protein-coding (eight genes) and tRNA genes (14 genes), and both rRNA genes (rRNA-L and rRNA-S). Missing from our mitogenome were the protein-coding genes *atp*6, *atp*8, *cox*3, *nad*3, and *nad*5, and the tRNA genes, tRNA-Ala, tRNA-Arg, tRNA-Asn, tRNA-Glu, tRNA-Gly, tRNA-His, tRNA-Phe, tRNA-Phe, tRNA-Trp, tRNA-Tyr, and a second copy of tRNA-Ser.

### Differential gene expression

We compared the transcriptomes of resistant (Keith and Temora) and susceptible (Kellerberrin and Eudunda) strains to identify candidate over-expressed genes linked to resistance. Three major patterns emerged from these transcriptomic analyses. First, the resistant Keith and Temora strains exhibited highly similar gene expression patterns. Second, gene expression patterns between the resistant strains were clearly distinct from those of the two susceptible Kellerberrin and Eudunda strains. Third, the Kellerberrin and Eudunda strains exhibited distinct gene expression patterns relative to each other. These patterns were evident in the clear separation of resistant from susceptible strains on PC axis 1 (41.89% of variation), and in the separation of the susceptible strains from each other on PC axis 2 (21.94% of variation) (Figure 1b).

Our primary goal for the transcriptome analyses was to identify genes that were over- expressed in the resistant strains relative to the susceptible strains. Because resistant and susceptible strains represented distinct clonal genetic background (see below), interpreting expressional differences requires care to not confound basal genetic differences with insecticide adaptation. However, as a first step, we used the transcriptome data to identify over-expressed detoxification genes as potential candidate insecticide resistance genes for further experimental validation.

After controlling for false positives, we identified 198 candidate genes that were consistently over-expressed in all four pairwise combinations of resistant versus susceptible strains (Supplementary Figure S3; Appendix 1). GO (gene ontology) enrichment analysis did not find any terms that were over-represented in these top over- expressed genes. Among the top over-expressed genes, seven were associated with potential detoxification functions: an E4-like esterase (gene ID g15286 with a 201-fold change); an ABCB transporter (gene ID g18463 with 4-fold change); two cytochrome P450s (gene IDs g15285 and g11751, with 6-fold change and 13-fold change, respectively); and three UDP-glycosyltransferases (gene IDs g2017, g14292, and g6298 with 4-fold, 3-fold, and 5-fold change, respectively).

Of greatest interest was the E4-like esterase gene for two reasons. Firstly, this was clearly top over-expressed gene in all pairwise comparisons between either the Temora or Keith strains and the susceptible Kellerberrin and Eudunda strains. Secondly, the protein sequence of this *A. kondoi* E4-like esterase showed ∼80% similarity to the *M. persicae* E4/FE4 esterase, which is known to contribute to organophosphate, carbamate and pyrethroid resistance in that species through over-expression that is linked to increased copy number (Field and Devonshire 1998; Field and Foster 2002). Similar esterases have also been shown to be over-expressed in other resistant aphid species (Ono et al. 1999; Müller et al. 2023). This E4-like esterase therefore became the focus of further experimental investigations.

### Clonal genetic background

We inferred evolutionary histories among clonal strains by calculating genetic distances using genome-wide biallelic SNPs. For the Keith, Temora, Kellerberrin and Eudunda strains, SNPs were derived from transcriptomes. For the Bangham, Coombe, Moulamein, and Jung strains, SNPs were derived from whole-genome sequencing data. Our comparison of genetic distances among all core eight strains indicated that the resistant strains formed a monophyletic lineage that was distinct from all susceptible strains regardless of geographic origin (Figure 1c).

Within the resistant and susceptible lineages, genetic differences were comparable across strains for 4,161 biallelic SNP loci. Comparing across resistant strains, the mean allelic differences ranged from 9 to 84.75 alleles, with a mean of 60.31 alleles. Comparing across susceptible strains, the mean allelic differences ranged from 9 to 141.25 alleles, with a mean of 84.82 alleles. When comparing within strains, the mean allelic differences for the four strains with transcriptomic data (Eudunda, Kellerberrin, Temora and Keith) ranged from 11.62 to 17.88 alleles, with a mean of 13.97 alleles. We could not provide the same within-strain assessment for the four strains with whole-genome data (Bangham, Coombe, Jung, Moulamein) because we only sequenced one pooled sample per strain. Nevertheless, across the resistant and susceptible lineages, allelic differences between strains ranged from 3,657 to 3,740 alleles, with a mean of 3,699 alleles. Hence, differences *within* broader resistant vs susceptible clonal genetic backgrounds were dramatically reduced compared to differences *between* them.

We noticed that samples genotyped with whole-genome sequencing were always nearest neighbours in our genetic distance trees and tended to sit paraphyletic to samples genotyped with transcriptome sequencing. It is known that allele-specific expression, splicing, or editing can cause issues with heterozygote detection from transcriptome data (Castel et al. 2015; De Wit et al. 2015). However, in our study, there were no clear reductions in SNP-level heterozygosity of transcriptome sequenced samples relative to whole-genome sequenced samples (Supplementary Table S2). If technical differences were at play, then a few key loci may have caused clustering of samples by the sequencing approach, but this pattern was not evident in the results.

We further examined the genetic divide between resistant vs susceptible *A. kondoi* by genotyping mitochondrial SNP loci. In principle, mitochondrial markers should not be affected by allele-specific expression because there is only a single allele within a clonal strain. We mapped our transcriptome and whole-genome sequencing data to our mitogenome assembly and identified candidate sites that were fixed between the resistant and susceptible strains. We then developed sets of primers to amplify focal regions in the *cytb*, *cox*1, and *cox*2 genes, each of which contained a diagnostic SNP for resistance status. These primers were used to genotype individuals from additional clonal strains (not included in our genomic datasets) via Sanger sequencing of PCR amplicons. Combining all mitochondrial data, clear genotypic differences were observed between resistant vs susceptible lineages, irrespective of sequencing approach, emphasising a strict association between the genetic background and the resistance phenotype (Figure 2, left panel).

**Figure 2.**
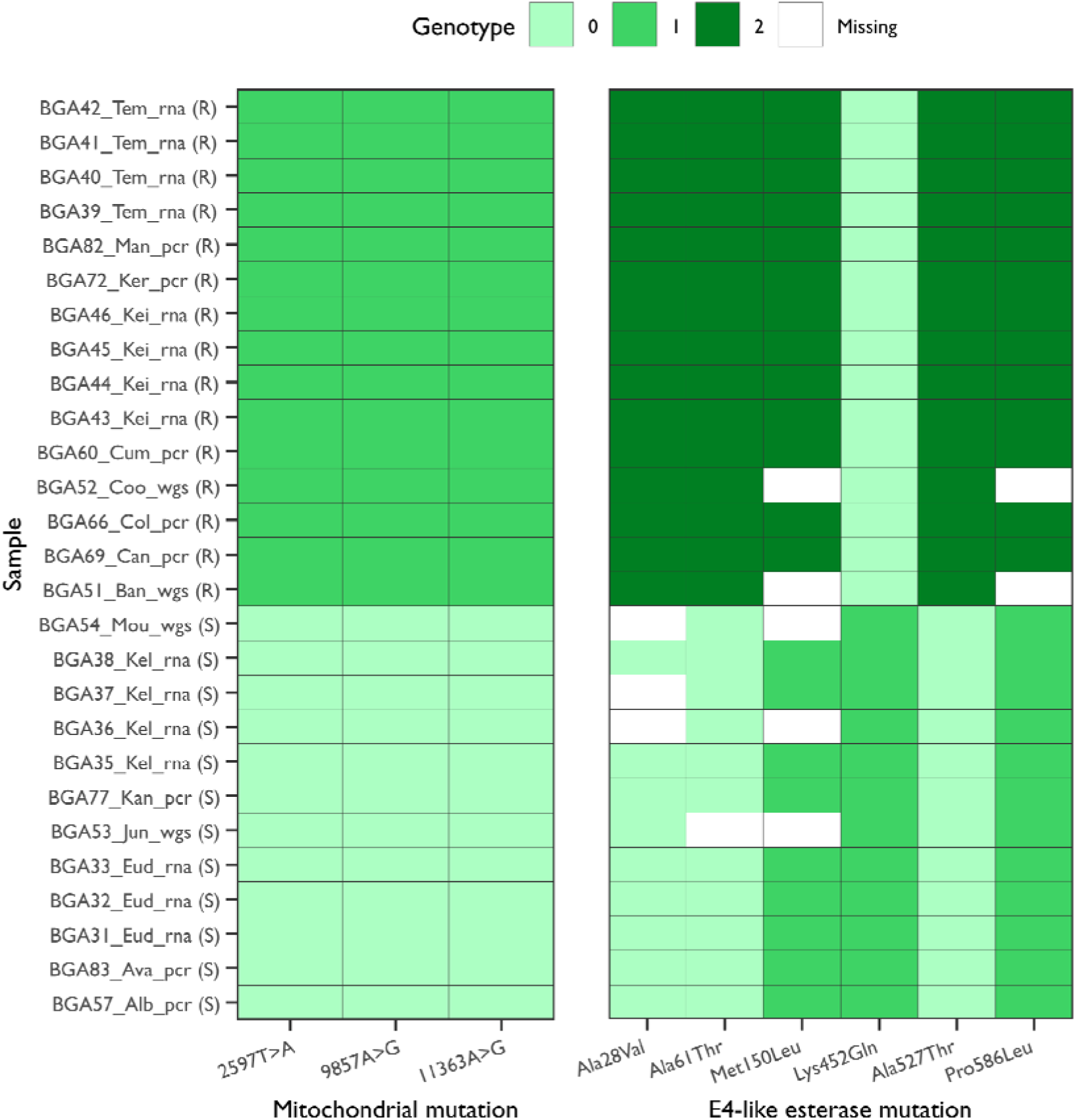
Diagnostic mutations in mitochondrial genes and the candidate E4-like esterase gene. Mutant (non-reference) alleles are arranged along the x-axis. Samples are arranged on the y-axis, labelled as ‘ID_Strain_Method (Status)’: ‘ID’ is a unique sample identifier; ‘Strain’ is strains name; ‘Method’ is one of ‘rna’ for transcriptome sequencing, ‘wgs’ for whole-genome sequencing, and ‘pcr’ for Sanger sequenced PCR amplicons; and ‘Status’ is one of ‘S’ for susceptible, or ‘R’ for resistant. Cell colours indicate the observed genotype (see legend).

### E4-like esterase genomic copy number, expression, and sequence variation

The identified *A. kondoi* E4-like esterase (a type B carboxylesterase) was an intriguing candidate for further investigation given its homology to esterases that confer insecticide resistance in other aphids. We used qPCR to validate the over-expression of the E4-like esterase in resistant *A. kondoi* and test whether this over-expression was achieved through increased copy number of this gene, as seen in *M. persicae* (Field and Devonshire 1998; Field and Foster 2002). A custom set of qPCR primers was designed for the candidate E4-like esterase as our focal gene to be compared against actin as the reference gene – the efficiency of these primers was tested before conducting our experiments (Supplementary Figure S4).

We conducted two experiments that compared relative amounts of RNA vs DNA within and between resistant vs susceptible strains: in Experiment 1, samples were comprised of RNA and DNA extracted from single aphids, and in Experiment 2, samples were comprised of two pooled aphids. As expected, RNA (cDNA) samples exhibited –ΔC_P_ values that were generally positive for resistant strains and negative for susceptible strains (Figure 3), supporting the hypothesis of over-expression in the resistant strains.

**Figure 3.**
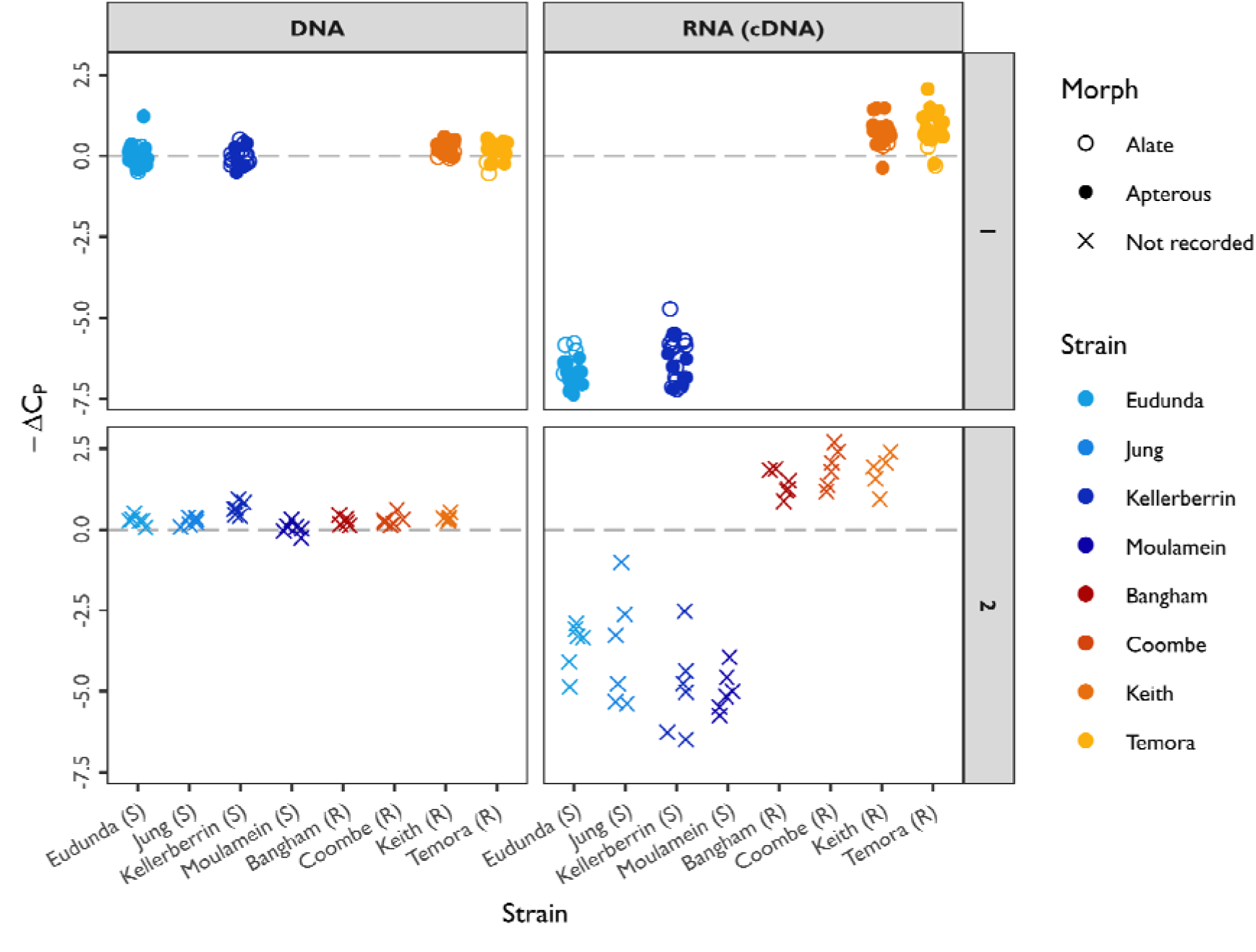
Among-strain comparisons of candidate E4-like esterase relative genomic copy number and expression in 12-day-old *Acyrthosiphon kondoi* measured with qPCR. Clonal strains are arranged on the x-axis. The –ΔC_P_ is on the y-axis, which represents the inverse of the ΔC_P_, where the ΔC_P_ = E4-like esterase C_P_ – actin C_P_. A positive –ΔC_P_ therefore indicates a higher amplification of the E4-like esterase (more molecules), and a negative –ΔC_P_ indicates a lower amplification of the E4-like esterase (fewer molecules), relative to the reference actin gene. Panels contain results from different experiments (rows: Experiment 1 and 2) and molecule types (columns: DNA and RNA/cDNA). The dashed grey line demarcates a –ΔC_P_ of 0, that is, no difference in the amplification of the E4-like esterase and actin genes. Points are coloured by strain, with shapes indicating the aphid morph (see legend). In Experiment 1: points represent individual-level measurements for the susceptible strains Eudunda and Kellerberrin, and the resistant strains Keith and Temora (morph type is indicated by shape, see legend). In Experiment 2: points represent pooled measurements (two aphids per sample) for the susceptible strains Eudunda, Jung, Kellerberrin, and Moulamein, and the resistant strains Bangham, Coombe, and Keith (morph type not recorded).

The relative fold-change estimated between resistant and susceptible strains using the 2^-ΔΔCP^method (Livak and Schmittgen 2001) (Supplementary Figure S5 and Supplementary Table S3) showed that in Experiment 1, the median fold-change ranged from 127- to 184-fold (mean of 155-fold), and in Experiment 2, the median fold-change ranged from 29- to 124-fold (mean of 75-fold). In contrast, the –ΔC_P_ values for DNA samples were ∼0 across strains (Figure 3), suggesting that both genes are present at equivalent genomic copy numbers and are amplified at the same rate in qPCR, irrespective of resistance status. This therefore does not support the hypothesis that over-expression of the E4-like esterase is achieved through increased copy number, so some other regulatory mechanism must therefore be at play.

Closer examination of sequence variation within the E4-like esterase revealed distinct genotypic differences between resistance and susceptible *A. kondoi* strain, irrespective of whether they were genotyped with transcriptome, whole-genome, or Sanger sequencing. A total of five non-synonymous mutations were observed in the candidate E4-like esterase: Ala28Val, Ala61Thr, Met150Leu, Ala527Thr, and Pro586Leu. Resistant strains were homozygous for the alternate allele at codons 28, 61, 150, 527 and 586, and homozygous for the reference allele at codon 452 (Figure 2, right panel). Susceptible strains had identical genotypes, which comprised homozygous reference alleles for codons 28, 61 and 527, and heterozygous genotypes at codons 150 and 586 (Figure 2, right panel).

To assess the possible functional effects of these non-synonymous mutations, we aligned the *A. kondoi* E4-like esterase against the *M. persicae* E4 esterase homolog and the *Torpedo californica* acetylcholinesterase as a model for type B carboxylesterases (Figure 4). We also produced a structural model of the *A. kondoi* E4-like esterase, which was of overall good quality (Figure 5): the ERRAT score was 93.91, and the Ramanchandran plot value was 86%, with low-confidence regions occurring in the disordered signal peptide sequence toward the N-terminus (Supplementary Figure S6). Our annotation of the multi-sequence alignment and structural model suggested that none of the non-synonymous mutations occur in, or near, the active site or interfere with the amino acids forming the catalytic triad involved in substrate hydrolysis (Ser221, Glu348, His461, *T. californica* numbering). Five of the non-synonymous mutations are located close to the surface of the protein, and mostly on the opposite side to the active site cleft (Figure 5); the exception was the Ala28Val mutation, located in the signal peptide. It is therefore unlikely that the observed non-synonymous mutations would dramatically alter the substrate profile of the enzyme.

**Figure 4.**
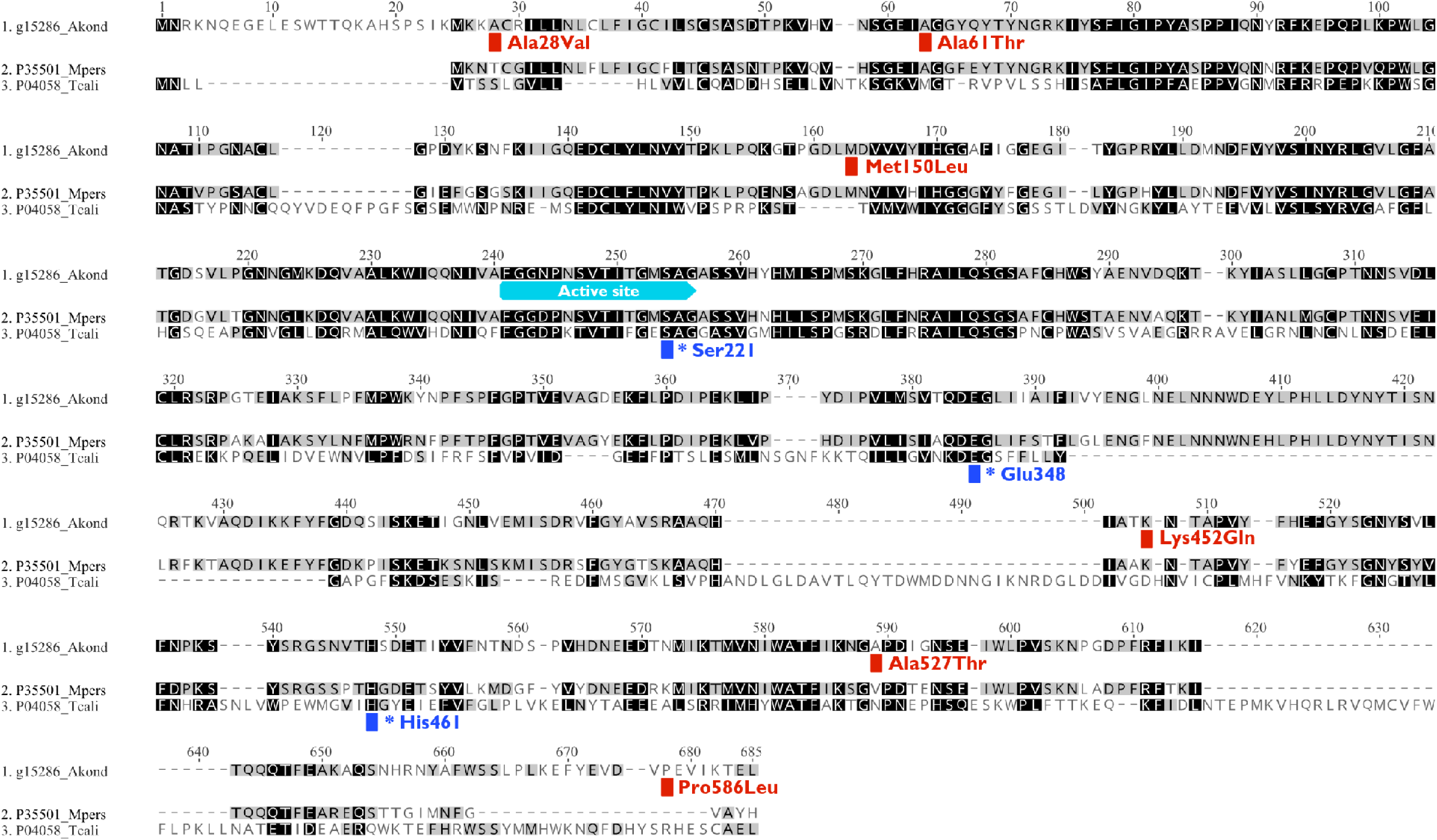
Protein alignment of the *Acrythosiphon kondoi* E4-like esterase to other type B carboxylesterases. The *A. kondoi* E4-like esterase (1) is compared against its homolog *Myzus persicae* E4 esterase (2), as well as the *Torpedo californica* acetylcholinesterase (3) as a classic example of type B carboxylesterase. Amino acids are shaded by their level of conservatism (darker shades are more conserved). Red annotations in the *A. kondoi* E4-like esterase sequence indicate positions where non-synonymous mutations are found (*A. kondoi* numbering). Cyan annotations indicate positions of the active site in the *A. kondoi* E4-like esterase sequence. Blue annotations mark the amino acids forming the catalytic triad in the *T. californica* acetylcholinesterase sequence (indicated with an asterisk, *T. californica* numbering). (See also Figure 5)

**Figure 5.**
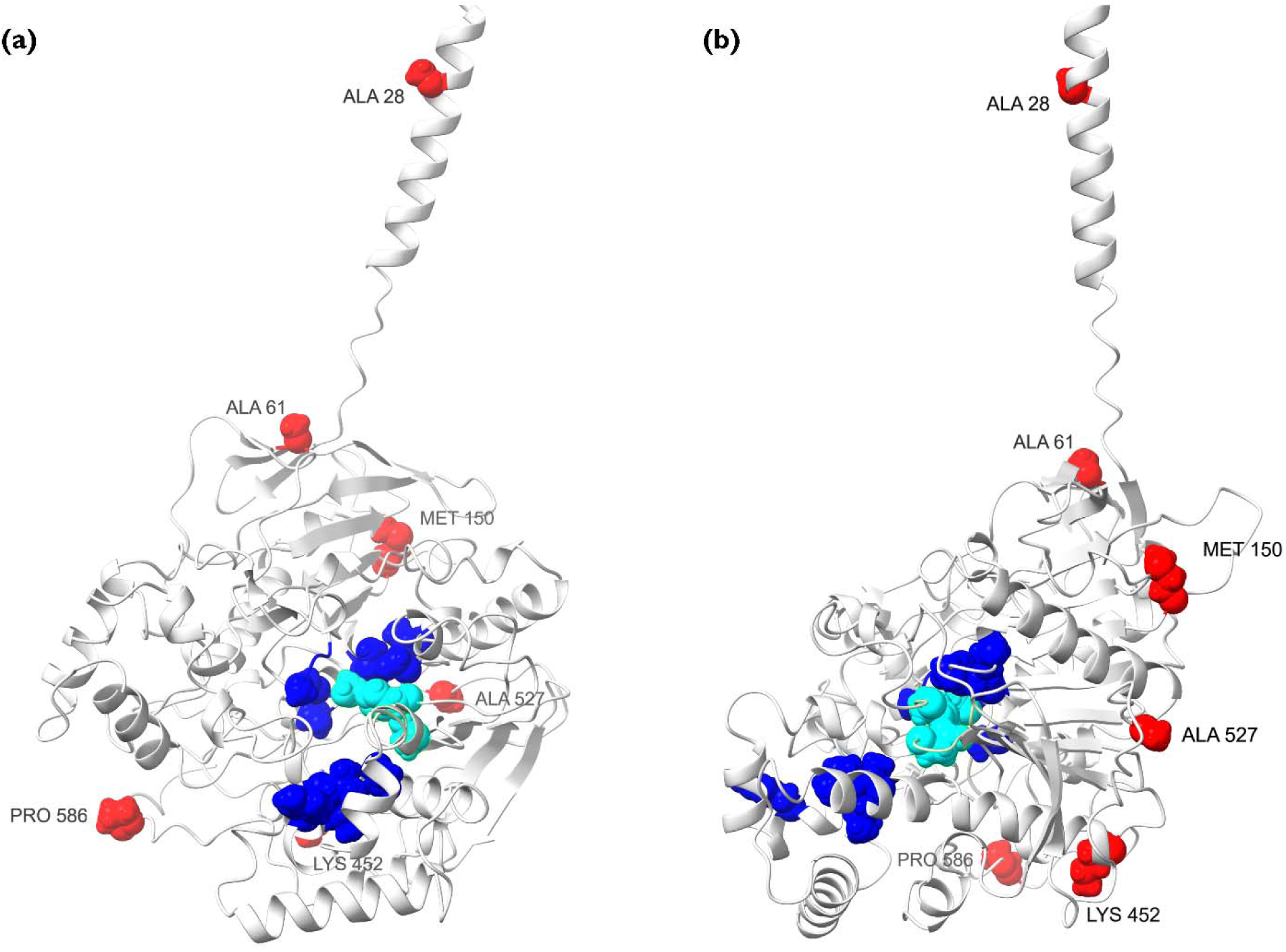
Structural modelling of the *A. kondoi* E4-like esterase protein. (a) Projection of the protein structure with the active site facing forward. The catalytic triad is highlighted in cyan, while the amino acids forming part of the substrate pocket are highlighted in blue. The positions of six non-synonymous mutations are highlighted in red and are labelled by their reference amino acid. (b) The same structure viewed from the side, showing that the positions of the non-synonymous mutations are located close to the surface on the opposite side of the active site. The exception is the Ale28 position, which is located on the signal peptide. (See also Figure 4)

### E4-like esterase transgenesis

Experiments using transgenic *Drosophila melanogaster* were used to functionally validate the role of the *A. kondoi* E4-like esterase in resistance. We successfully established three transgenic strains of *Drosophila* expressing different version of the E4-like esterase, which we called, H1, H2, and H3. H1 was identical to the susceptible reference genome sequence (Figure 2, right panel). H2 contained the mutations Met150Leu and Pro586 Leu, which are heterozygotes in susceptible *A. kondoi* and homozygous in resistant *A. kondoi* (Figure 2, right panel). H3 contained three mutations, Ala28Val, Ala61Thr, and Ala527Thr, absent in the susceptible *A. kondoi*, and homozygous in the resistant *A. kondoi* (Figure 2, right panel).

Using insecticide bioassays, we demonstrated that ectopic expression of *A. kondoi* E4- like esterase in *D. melanogaster* conferred significant resistance to three insecticides, relative to a control strain of the same genetic background but lacking this gene (Table 2 and Figure 6). The level of resistance conferred by E4-like esterase in *D. melanogaster* was modest, ranging from a 1.8-fold to 4.9-fold increase depending on the insecticide and haplotype tested (Supplementary Table S4). For logistical reasons, we were unable to compare all haplotypes in the same experiment, which limited a direct three-way comparison (see Materials and Methods). However, we could directly compare H1 (susceptible-like) and H3 (resistant-like), which were run in the same experiment.

**Figure 6.**
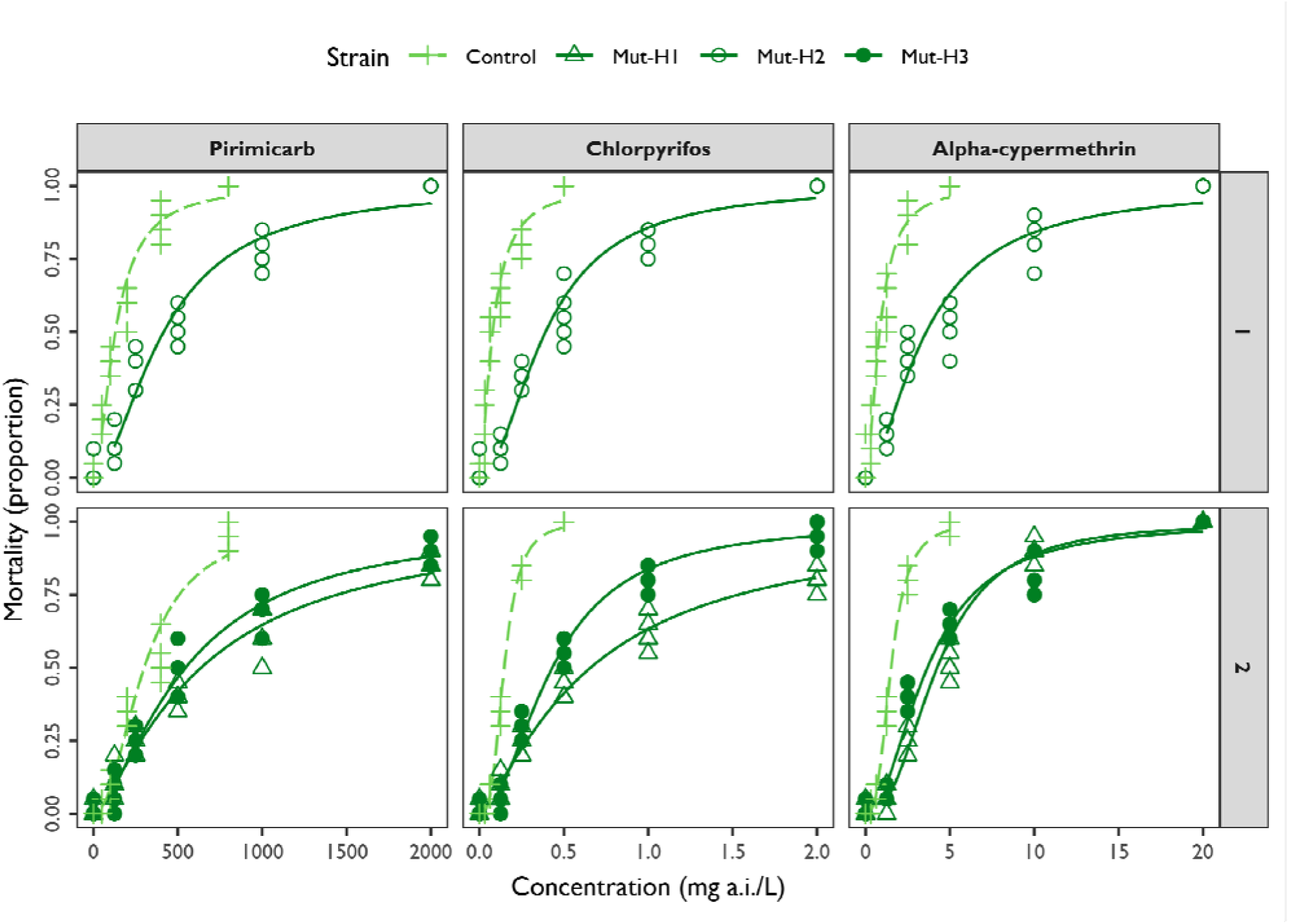
Dose-responses generated from insecticide bioassays on transgenic *Drosophila melanogaster* carrying our candidate E4-like esterase from *Acythosiphon kondoi*. The x-axis represents the concentration of insecticide, and the y-axis represents the proportion of mortality. Points represent individual replicates with predicted dose response curves (line typeand colour depicts strains, see legend). Panels contain data for different experiments (rows) and insecticides tested (columns) (see also Table 2).

The H1 and H3 haplotypes had diametrically opposed genotypes at codons 28, 61 and 527 between resistant and susceptible *A. kondoi* strains (Figure 2, right panel). We found that H1 exhibited a higher LD50 relative to H3 across all insecticides, but this was only significant for chlorpyrifos (where confidence intervals did not overlap) (Table 2). This pattern is counter to what we might expect if mutations in the resistant-like H3 haplotype were important for the detoxification of insecticides in *A. kondoi*. These results further support the mechanism of resistance of the *A. kondoi* E4-like esterase involving over-expression and not protein-coding changes in this gene.

### Target-site mutations

Insecticides typically exert their toxicity by targeting specific genes. Organophosphates and carbamates target the acetylcholinesterase (*ace*), and pyrethroids target the para- like voltage-gated sodium channel (*para*). We therefore examined SNPs in the exonic regions of these genes to assess whether any might provide target-site resistance to their associated insecticides. Our genome contained three genes with *ace*/*ace*-like annotations (g46, g11959, and g15835), but none exhibited non-synonymous mutations. Our genome contained two *para* gene annotations (g11612 and g11614), but we only observed three non-synonymous mutations in one *para* gene, g11614, which included Thr104(1201)Ala, His359(1450)Asn, and His359(1450)Gln (note that codon numbers in parentheses are the position from canonical *para* reference sequence from *Musca domestica*) (Supplementary Figure S7). Comparison of our two *A. kondoi para* genes with the pest aphids *A. pisum* and *M. persicae* indicated that gene g11614 is the *para* subunit 2, whereas gene g11612 is the *para* subunit 1 (Supplementary Figure S7). The structure of these two genes in the genome of *A. kondoi* is very similar to that in *A. pisum* and to that described in *M. persicae* (Amey et al. 2015): both genes occur in a proximate genomic region but are on different strands (Supplementary Figure S8). While susceptible strains are homozygous for the reference mutations in g11614 *para* subunit 2, all resistant strains are heterozygous (Supplementary Figure S9). These mutations have not been observed in other pyrethroid-resistant arthropods (Dong et al. 2014). Moreover, documented pyrethroid target-site mutations in aphids appear to evolve in subunit 1 (Amey et al. 2015; Zuo et al. 2016; Valmorbida et al. 2022), not subunit 2. We thus take the conservative interpretation that mutations observed in our study are unlikely to be involved in pyrethroid resistance, and that there is no evidence for target-site resistance in *A. kondoi* generally.

## Discussion

The evolutionary origins and repeatability of adaptation have long intrigued evolutionary biologists (Cohan and Hoffmann 1989; Korona et al. 1994; Colosimo et al. 2005; Roda et al. 2013; Tepolt et al. 2021). Here, we developed the first reference nuclear genome and partial mitogenome for the aphid, *Acyrthosiphon kondoi,* an economically important global pest. We used these new genomic resources to study the evolutionary origins and molecular mechanisms of insecticide resistance in its invasive Australian range.

Insecticides are the major way in which *A. kondoi* is managed in Australian agriculture, with organophosphates, carbamates, and synthetic pyrethroids widely used in alfalfa growing regions. This prolonged selection pressure has led to the evolution of resistance, first detected after farmers reported chemical control failures in a small number of field populations in 2018 (Chirgwin et al. 2023). Although not registered to control *A. kondoi* in most cropping situations and thought to be infrequently used against this species in Australia, resistance was also detected to multiple synthetic pyrethroids, namely alpha-cypermethrin and gamma-cyhalothrin (Chirgwin et al. 2023). Insecticide resistance is a trait that is often broadly beneficial in pest organisms due to the use of agrichemicals across large areas (Chevillon et al. 1999; Endersby-Harshman et al. 2020; Zhang et al. 2023). Moreover, clonality in asexual aphids can provide the added benefit of rapidly spreading resistant lineages in local populations and facilitating recovery following major selection events (Muller 1932; Crow and Kimura 1965; Vorburger 2006; Loxdale 2008; Loxdale et al. 2010).

Our genomic investigation shows that all resistant clonal strains shared a single evolutionary origin and are highly differentiated from all other susceptible clonal strains examined here. The resistant lineage exhibits resistances to all three insecticide classes (organophosphates, carbamates, and pyrethroids) used to control it in Australia, undoubtedly providing a selective advantage for the aphid within the context of Australia’s alfalfa and legume crops where there are limited chemical control options.

Resistant strains analysed in this study were collected hundreds of kilometres apart. This geographic spread of a single resistant clonal lineage of *A. kondoi* is characteristic of superclones in other aphid species. In Australian populations of *Myzus persicae*, more than 300 unique clones have been characterised using microsatellites, yet there is very limited genetic variation among aphids collected from agricultural fields from different areas due to selective sweeps of resistant superclones (Umina et al. 2014; de Little and Umina 2017; Thia, Zhan, et al. 2023). Similarly, a resistant clone of the English grain aphid, *Sitobion avenae*, has displaced sexually reproducing lineages in the UK, resulting in reduced genetic variation in the field (Morales-Hojas et al. 2020). An important question arising from this work is how long resistance will persist and (or) continue to spread in the field. This will depend on whether selection pressures can be relaxed to reduce resistance by exploiting costs of resistance, as observed in other arthropod pests (Foster et al. 1999; Bass et al. 2014; Zhang et al. 2015; Maino et al. 2018) However, if the costs of resistance are minimal, resistant *A. kondoi* clones may continue to persist even if insecticide usage is reduced.

The most common mechanisms of evolved insecticide resistance involve target-site changes and the evolution of detoxification mechanisms. In this work, we did not find any clear evidence that resistance in *A. kondoi* is linked to target-site mechanisms. We identified no non-synonymous mutations in any annotated *ace* or *ace*-like genes. The only three non-synonymous mutations detected were in the g11614 *para* gene, but these are not associated with resistance in other arthropods. Instead, our data provide strong evidence that insecticide resistance in *A. kondoi* has evolved through detoxification mechanisms. Using transcriptomic analyses, we identified a type B carboxylesterase, an E4-like esterase, as a candidate gene for resistance. Relative to the susceptible strains, resistant strains exhibited over-expression of this E4-like esterase in the range of 29-fold to 184-fold in qPCR assays and >200-fold in transcriptomic analyses.

The *A. kondoi* E4-like esterase is an intriguing candidate for resistance given is its similarity to the E4/FE4 esterases in *M. persicae,* which are known to be major detoxifiers of organophosphate, carbamate, pyrethroid insecticides (Field and Devonshire 1998; Field and Foster 2002). Homologs of this esterase family have also been identified in other aphid species with evolved resistance. In the wheat aphid, *Schizaphis graminum*, an E4/FE4 esterase homolog has been linked to organophophate resistance (Ono et al. 1999). In the pea aphid, *A. pisum*, two recently described pyrethroid resistant strains appear to over-express an FE4 esterase homolog, although it is still unclear whether this esterase contributes to resistance (Müller et al. 2023). In both *M. persicae* and *S. graminum*, over-expression of these esterases is achieved through gene amplification, with additional regulation through methylation patterns (Hick et al. 1996; Field et al. 1999; Ono et al. 1999; Field 2000). Whilst there appears to be an intriguing macroevolutionary parallelism at the gene family level between *M. persicae*, *S. graminum* and *A. kondoi*, the regulatory mechanism in *A. kondoi* appears to be different in that it is not associated with gene amplification. Exploring the role of promoter sequence or epigenetic variation is a possible avenue for future research.

Functional analysis of the *A. kondoi* E4-like esterase demonstrated that this enzyme is a general detoxifier of insecticides. In our bioassays, transgenic *Drosophila* ectopically expressing the *A. kondoi* E4-like esterase were significantly more resistant to alpha-cypermethrin (a pyrethroid), chlorpyrifos (an organophosphate), and pirimicarb (a carbamate). This highlights the ability for this esterase to provide cross-resistance across different chemical groups, effectively acting as a gene with pleiotropic effects. Our current data does not support the role of protein-coding mutations in the E4-like esterase contributing to resistance in *A. kondoi*. None of the six non-synonymous mutations observed in this study occurred in the active site or were proximal to the catalytic triad (Ser238, Glu363, His487, *A. kondoi* numbering) that mediates hydrolysis (Cygler et al. 1993; Hatfield et al. 2016). Although it is possible that mutations distal to an active site can impact catalytic activity (Amichot et al. 2004), we did not see differences in our transgenic *Drosophila* experiments. The resistant-like H3 *Drosophila* strain carrying Ala28Val, Ala61Thr, and Ala527Thr, did not exhibit greater resistance than the susceptible-like H1 strain carrying no mutations. A caveat of this work was that we were unable to compare the H2 *Drosophila* strain carrying the Met150Leu and Pro586Leu mutations to the H1 and H3 strain, and we were also unable to generate a strain containing the Lys452Gln mutation. However, the Met150Leu and Pro586Leu are heterozygous in susceptible *A. kondoi*, and resistant *A. kondoi* are homozygous for the reference Lys452 allele. Hence, Met150Leu, Lys452Gln and Pro586Leu are weaker candidates for resistance. There is scope for additional experimental work and protein modelling to further test the hypothesis that protein-coding differences in the E4-like esterase contribute to resistant phenotypes. But for now, we suggest that over- expression is the main mechanism through which this gene contributes to resistance in *A. kondoi*.

We note that whilst resistance in our mutant *Drosophila* strains was significant, it was also modest, with resistance ratios ranging from 1.81 to 4.88, relative to the non-mutant *Drosophila* strain. This in a sense mirrors the modest to moderate resistance ratios observed in *A. kondoi*, which has been reported in the range of 6.80 to 20.26 depending on the resistant strain and insecticide tested (Chirgwin et al. 2023). Although our data supports the E4-like esterase as an important gene contributing to insecticide resistance, there are likely other mechanisms that contribute to the phenotypic variance in resistance between the *A. kondoi* genetic lineages. In our differential expression analyses, for example, cytochrome P450s, UDP-glucuronosyltransferases, and an ABCB transporter were also among the top over-expressed genes. Exploring the roles of these other genes will be the subject for future work.

Our reference nuclear and (partial) mitochondrial genomes for *A. kondoi,* together with the detailed transcriptomic analysis, provides an important contribution to the limited genomic resources available for this economically significant aphid species. The high BUSCO score (97.5%) for our nuclear genome indicates that this is a high-quality assembly, with 615 contigs and a total length of 443.8 Mb. Although we were only able to assembly a partial mitogenome, this still represents the longest mitochondrial sequences from *A. kondoi* to date, capturing most of the protein coding and tRNA genes, and both the rRNA genes. These genomic resources allowed us to uncover key insights into the evolution and genetic mechanisms underlying insecticide resistance in *A. kondoi* and assisted in developing diagnostic markers for the resistant genetic lineage. Such diagnostic markers will facilitate the collection of spatiotemporal data on resistance, providing insights into the dynamics of the resistant clonal lineage.

## Concluding remarks

In this work, we disentangled the hypotheses of parallel evolution vs the spread of insecticide resistance in *A. kondoi*, an important agricultural pest. Through genomic and transcriptomic analyses, we revealed that a major component of insecticide resistance in *A. kondoi* is due to detoxification by an over-expressed E4-like esterase. This esterase shares homology with other resistance-conferring esterases in different aphid species. Our work also highlights the rapid spread of a single resistant superclonal lineage, reflecting the influence of human-induced selection pressures on pest populations. While our findings suggest a parallel macroevolutionary response to insecticide exposure in *A. kondoi* and other aphid species, they also suggest a unique regulatory mechanism of the *A. kondoi* E4-like esterase, warranting further investigation. The cross-resistance provided by the E4-like esterase (and other potential detoxification genes) poses a serious challenge to management because multiple insecticide classes may reinforce selection on the same genetic mechanism(s). Given the rapid spread of resistance in Australia, there is a need to develop alternative management strategies for *A. kondoi*, including expanding chemical control options and enhancing the use of biological control.

## Materials and Methods

### Aphid strains

Eight clonal isofemale strains formed the focal strains for our experiments. These strains were generated with artificial selection using leaf-dip insecticide bioassays (Umina et al. 2014). Nymphs from Keith, Temora, Bangham and Coombe were exposed to alfalfa leaves dipped in 15 mg a.i./L chlorpyrifos (Lorsban 500EC, Corteva) for 72 hours. This concentration was previously shown to effectively separate resistant aphids from insecticide-susceptible individuals (Chirgwin et al. 2023). The chlorpyrifos solution was prepared by diluting 15 mg a.i./L in distilled water. Leaves were dipped in the insecticide solution for 10 seconds and then plated onto 50 mm Petri dishes containing 1% agar.

Three dishes were prepared for each strain, each containing three large alfalfa leaves and 10 three-day-old nymphs. After 72 hours, surviving aphids were separated and individually plated on 30 mm Petri dishes with a single untreated alfalfa leaf on agar.

Approximately eight days later, individuals that survived and produced nymphs were retained in separate Petri dishes. After three generations, the healthiest surviving replicate was picked to start an isofemale strain. Such strains were also generated for the susceptible Kellerberrin, Eudunda, Jung, and Moulamein strains, but these were not exposed to artificial selection. A follow up bioassay using the same discriminating dose of chlorpyrifos (15 mg a.i./L) confirmed the resistance status of these isofemale strains (Supplementary Table S1).

We later sampled additional field strains to broaden the geographic scope of our study, which were from Albany, Avalon, Kaniva, Canowindra, Coolah, Cummins, Kerang, and Manoora. Whilst we know the resistance status of these field strains (Evatt Chirgwin, unpublished data), we did not subject them to artificial selection and screening procedures in the same way as for our core experimental strains. Single individuals were sampled from these field strains for genetic analysis (see below). Genomic DNA from these individuals was extracted using a Qiagen EZ2 Tissue Kit.

### Genome assembly and annotation

#### Nuclear genome

We developed the first reference for the *A. kondoi* nuclear genome to investigate the genomic basis of insecticide resistance. The susceptible (laboratory) Kellerberrin strain was used for the assembly. We performed two high molecular weight DNA extractions from 22 and 24 pooled adult aphids, respectively, using the Nanobind Tissue Kit following a modified ‘UHMW DNA Extraction – Dounce Homogenizer’ protocol. However, we replaced the Dounce Homogeniser with a simpler homogenisation step, crushing aphids in a 1.5 mL Eppendorf tube with a pestle chilled on ice. The tissue homogenate, lysis buffer, and proteinase K were incubated overnight at 50°C. All other steps were as per the protocol. The DNA was sent to Novogene (Singapore) for library preparation, long-read sequencing using the Oxford Nanopore Technology (ONT) PromethION platform, and short-read Illumina paired end (150 bp) sequencing on the NovaSeq platform.

We also sequenced a transcriptome from 10 pooled 14-day-old apterous aphids from the Kellerberrin strain. To synchronize aphids, 15 adults were placed onto a 100 mm Petri dish with alfalfa leaves in agar. These aphids were given an initial 24 hours to settle, after which any offspring produced were removed. After a further 24 hours, the adults were removed and any offspring produced were left behind on plates. Offspring were re- plated every 3 to 4 days, and any showing signs of developing into alates were removed. After 14 days, apterous aphids were pooled, frozen in liquid nitrogen, and sent to AGRF (Australian Genome Research Facility, Melbourne) for RNA extraction, library preparation, and short-read Illumina sequencing (150 bp) on the NovaSeq platform. In total, we obtained 7,076,013 ONT long-reads, 191,891,532 paired reads from the Illumina whole- genome sequencing, and 11,532,979 paired reads from Illumina transcriptome sequencing. Illumina reads were quality trimmed using FASTP v0.23.2 (Chen et al. 2018).

Whole-genome short reads were used to estimate the genome size using a combination of the KMC v3.1.2 (Kokot et al. 2017) and GENOMESCOPE v2 (Vurture et al. 2017) programs. Our preliminary assessment of genome size for *A. kondoi* was ∼452 Mb, with a genomic heterozygosity rate of ∼0.87%, based on GENOMESCOPE. High-quality ONT long-reads were assembled with FLYE v2.9 (Kolmogorov et al. 2019), followed by three iterative polishing steps with whole-genome short reads using PILON v1.23 (Walker et al. 2014).

ONT long reads were mapped back onto this assembly with MINIMAP2 v2.17 (Li 2016) and duplicated regions were purged using PURGE_HAPLOTIGS v1.0.4 (Roach et al. 2018).

Repetitive regions were annotated using RED v2018.09.10 (Girgis 2015). Repeats were annotated using REPEATMODELER v2.0.4 (Flynn et al. 2020).

To annotate genic regions, RNA short reads were mapped to the curated assembly with HISAT2 v2.1.0 (Kim et al. 2019). All short-read alignments were filtered for quality (Q20) and duplicate reads and sorted using SAMTOOLS v1.17 (Li et al. 2009; Danecek et al. 2021). Gene annotation was performed with the BRAKER v.2.1.6 pipeline. The output AUGUSTUS (Stanke et al. 2006) gene annotations were retained for downstream analysis. Parsing of annotations and extraction of the longest protein coding transcript was performed through a combination of GFFREAD v0.12.7 (Pertea and Pertea 2020), AGAT v1.1.0 (Dainat 2022), and SEQKIT v2.0.0 (Shen et al. 2016). We assigned function to gene annotations using a combination of BLASTP v0.23.2 (Altschul et al. 1990) and BLAST2GO (Conesa et al. 2005; Conesa and Götz 2008). We downloaded the entire ‘nr’ database on the 10 of June 2023. We used BLASTP to match the amino acid sequences of the longest transcripts against all arthropod sequences in the ‘nr’ database. Gene ontology (GO) terms were assigned using INTERPROSCAN v5.59-91.0 (Zdobnov and Apweiler 2001) and revaluated using BLAST2GO. Assembly statistics were generated using QUAST v5.1.0 (Gurevich et al. 2013) and the completeness of single copy ortholog genes was assessed using BUSCO v3.1.0 (Simão et al. 2015).

We further assessed our nuclear assembly for redundancy using a self-to-self alignment with MINIMAP2. We also assessed completeness by aligning our *A. kondoi* assembly against the *A. pisum* chromosome-level assembly (NCBI Accession GCF_005508785.2). We first used the program RAGTAG (Alonge et al. 2022) to generate pseudo- chromosomes for *A. kondoi* using the *A. pisum* reference genome as a guide. The pseudo-chromosomes were mapped back to the *A. pisum* reference genome with MINIMAP2. We visualised dotplots and synteny between the *A. kondoi* pseudo- chromosomes and *A. pisum* chromosomes with R’s PAFR v0.0.2 package (Winter 2020).

#### Mitochondrial genome

We assembled a partial mitogenome reference for *A. kondoi*. We first mapped Illumina-short-reads from the Kellerberrin strain to our nuclear reference genome using BOWTIE2 and extracted the non-mapped reads with SAMTOOLS. The non-mapped reads were used to assemble a mitogenome reference with the program MITOFINDER v1.4.2 (Allio et al. 2020), using the *A. pisum* mitogenome as a guide (GenBank Accession NC_011594.1). We did attempt to perform an assembly from ONT long-reads but found that these yielded poor assemblies relative to the short-reads (unpublished data).

### Differential gene expression

We examined patterns of differential expression among the Keith, Temora, Kellerberrin and Eudunda strains to identify genes contributing to insecticide resistance in *A. kondoi*. Transcriptomes were derived from pools of 12-day-old apterous aphids. As above, we plated 15 adults onto 100 mm Petri dishes containing alfalfa leaves in agar alfalfa, with a 24-hour settling and culling period, and offspring after 48 hours were removed. These offspring were replated onto 30 mm Petri dishes containing a single alfalfa leaf on agar at densities of five to eight aphids per dish. After 8 days, aphids were moved to a 50 mm Petri dish containing two alfalfa leaves on agar. On day 12, apterous aphids from each strain were removed and snap frozen in liquid nitrogen. Four pools of aphids were used, with each pool comprising approximately 20 to 25 aphids (approximately 10 to 15 mg of tissue). RNA extraction was performed by OzOmics (Melbourne, Australia) using a Monarch Total RNA Miniprep Kit (New England BioLabs). Library preparation was performed by AGRF (Melbourne) with 150 bp paired-end sequencing on their NovaSeq platform. We successfully derived transcriptomes from all pools except one pool from the Eudunda strain. Hence, *n* = 3 for Eudunda, and *n*= 4 for all other strains. For the successfully sequenced transcriptomes, we obtained 35,232,161 to 52,797,301 raw reads, with a mean of 40,452,782 reads.

Raw transcriptomic reads were trimmed using FASTP. Trimmed transcript reads were then mapped to our assembled reference genome with HISAT2. Mapped reads were filtered to retain high-quality mapped reads (MAPQ ≥ 30) and were sorted using SAMTOOLS. Expression was quantified using STRINGTIE v2.2.1, and the STRINGTIE’s PREPDE.PY script was used to summarise data into a matrix of transcript abundances for all samples. This transcript abundance matrix was imported into R for differential expression analysis among strains. Global patterns of gene expression were visualised by analysing all strains together with DESEQ2 v1.40.2 (Love et al. 2014) and performing a PCA on the transcript abundances.

Differentially expressed genes between resistant and susceptible strains were identified using a pairwise approach. This involved tests on all possible pairs of resistant and susceptible strains (four total combinations), which allowed us to identify genes that were on average differentially expressed irrespective of the strains compared. Differential expression was determined with functions from the packages DESEQ2 v1.40.2 (Love et al. 2014) and APEGLM v1.22.1 (Zhu et al. 2019) in R. Our pairwise models predicted expression as a function of ‘Strain’ as a categorical factor with two levels (one for each strain). Within each pair, genes were ranked based on their *p*-value. We also separately generated the top 500 over- and under-expressed genes per pair.

Next, we identified candidate genes for resistance. Because our study was primarily interested in identifying detoxification genes, which are typically over-expressed in resistant strains, we focused further analyses on the top over-expressed genes.

Candidate genes needed to satisfy two criteria: (1) they were in the top 500 genes in a pairwise comparison, and (2) they were found across all four pairwise comparisons.

Mean ranks were generated based on the mean *p*-value across pairs.

Tests for GO term enrichment were performed on significantly over-expressed genes. Any gene that was significantly over-expressed in all four pairwise comparisons was considered a true outlier, and their GO terms used in enrichment analyses. These analyses involved Fisher’s exact test in R with the FISHER.TEST function, and *p*-values adjusted for multiple testing with a Bonferroni correction.

### Clonal genetic background

#### Nuclear genome

We tested whether resistant strains of *A. kondoi* were derived from the same clonal genetic background. For these analyses, we combined transcriptomic sequencing data from the Keith, Temora, Kellerberrin and Eudunda strains (above) with additional whole-genome sequencing data from the Bangham, Coombe, Jung and Moulamein strains. Whole-genome data was obtained by extracting genomic DNA from 6 pooled adult aphids using a Monarch Genomic DNA Purification Kit (New England BioLabs), one pool per strain. Whole-genome library preparation was performed by Novogene (Hong Kong) with 150 bp paired-end sequencing on their NovaSeq platform. We obtained 36,680,245 to 41,806,538 raw reads, with a mean of 40,405,084 reads.

Like the transcript reads, whole-genome reads were trimmed with FASTP, mapped with HISAT2, and filtered, sorted and deduplicated with SAMTOOLS. We then called variants in exonic regions by combining the mapped transcript and whole-genome reads using BCFTOOLS (Li 2011; Danecek et al. 2021). We then reduced our dataset to only contain SNP (single nucleotide polymorphism) loci with quality scores ≥30 using BCFTOOLS. Variants were imported into R using GENOMALICIOUS v 0.7.8 (Thia and Riginos 2019) and further filtered to only include SNPs with depth ≥15 reads, 0% missing data, and biallelic sites. The final dataset contained 4,161 biallelic SNPs.

Pairwise genetic distances between samples were quantified as the number of allelic differences across all biallelic loci. The pairwise genetic distance matrix was used to derive a UPGMA clustering tree with the PHANGORN v2.12.1 package (Schliep 2011), with negative branch lengths adjusted using the BAT v2.9.6 package (Cardoso et al. 2015).

Bootstrapping was undertaken with the APE v 5.8 package (Paradis et al. 2004). The resulting tree was visualised with GGTREE v3.12.0 (Yu et al. 2017) and TREEIO v1.28.0 (Wang et al. 2020).

#### Mitochondrial genome

We analysed mitogenomic variation as supporting data for the clonal genetic background of our *A. kondoi* strains. We first mapped transcriptome and whole-genome sequencing reads for our core strains against our mitogenome reference with HISAT2 and filtered alignments with SAMTOOLS to remove reads with quality scores <30. Variants were called with BCFTOOLS and were filtered to remove those with quality scores <30. Variants were imported into R and analysed with a combination of DATA.TABLE (Dowle and Srinivasan 2019), TIDYVERSE (Wickham et al. 2019), and GENOMALICIOUS. From these variants, we identified three diagnostic mitochondrial SNPs that distinguished resistant and susceptible strains: positions 2,597 (in *cytb*), 9,857 (in *cox*1), and 11,363 (in *cox*2).

We developed primers to amplify mitogenomic regions containing the three diagnostic SNPs using GENEIOUS PRIME (Supplementary Table S5). These primers were used to genotype representative individuals from the field strains. PCR amplification was performed in 25 uL volume comprising 1.25 uL MgCl_2_ (50 mM), 2.5 uL ThermoPol standard buffer (Mg free), 1.25 uL each of forward and reverse primer (10 mM), 2 uL dNTPs (2.5 mM), 0.4 uL IMMOLASE Taq polymerase, and 1.5 uL template DNA. PCR conditions were denaturation at 95°C for 10 mins, 45 cycles of 94 °C for 1 min, 56 °C for 1 min and 72 °C for 1 min, with a final extension at 72 °C for 7 mins. Amplicons were sent to Macrogen (South Korea) for Sanger sequencing in both forward and reverse directions. Raw sequences were quality checked and formed into consensus sequences in GENEIOUS PRIME. We also used GENEIOUS PRIME to map consensus sequences to our mitogenome reference and these alignments were used to score genotypes at our diagnostic SNPs.

### E4-like esterase genomic copy number, expression, and sequence variation

#### Copy number and expression

Our top candidate over-expressed gene for resistance was an E4-like esterase, which was over-expressed in resistant strains relative to susceptible strains (see Results). We used qPCR to validate the expression levels of the E4-like esterase, and to test whether elevated genomic copy number was driving over- expression. We performed two separate experiments with aphids for qPCR being obtained using the same protocols as those for obtaining transcriptomes (see Section, ‘Differential gene expression’). In Experiment 1, we measured individual levels (one aphid per sample) of genomic copy number and expression for the Eudunda, Kellerberrin, Keith and Temora strains. These individual measures also allowed us to test for expression differences between morphs (winged alate versus non-winged apterous aphids). For each strain, we analysed 24 samples of individual aphids. In Experiment 2, we measured genomic copy number and expression in pooled samples (two aphids per sample) for the Jung, Moulamein, Bangham and Coombe, Eudunda, Kellerberrin, Keith. Aphids for both experiments were obtained using the same protocols as those for obtaining transcriptomes (see Section, ‘Differential gene expression’). The Temora strain was not included in Experiment 2 due to a lack of nymphs during the age-matching step. In Experiment 2, we analysed five to six samples (two aphids per sample) because our findings from Experiment 1 indicated that individual levels of expression were consistent within strains and that there were no differences among morphs (see Results).

We developed a workflow that allowed us to simultaneously extract both the DNA and RNA from the same sample. Aphid samples (single aphids, or a pool of two aphids) were frozen in liquid nitrogen in a 1.5 mL Eppendorf tube and stored for up to two weeks at – 80°C. These were homogenized on ice with a small plastic pestle. The used pestle was then placed into a second Eppendorf tube containing 150 µL of 5% Chelex buffer and proteinase K (20mg/mL, Meridian, CSA-01254; Cincinnati, OH, USA) and swirled in this buffer mix to suspend the residual aphid homogenate − this was used for DNA extraction. We followed the protocol in Yang et al. (2023) for Chelex DNA extraction.

Simultaneously, 200 µL of RNA protection reagent from the Monarch Total RNA Miniprep Kit (New England BioSciences, Notting Hill, VIC, Australia; #T2010) was added directly to the homogenized aphid tissue in the original sample tube; this was used for RNA extraction. We performed RNA extraction following the Monarch Total RNA Miniprep Kit protocol. Extracted RNA was converted into cDNA using the ThermoFisher Scientific High-Capacity cDNA Reverse Transcription Kit (Parkville, VIC, Australia; #4368814) as per manufacturer’s instructions.

We used qPCR to quantify the relative expression of the candidate E4-like esterase across clonal strains of *A. kondoi*. DNA extractions were diluted 1:3 and cDNA samples were diluted 1:10 with PCR water. A set of primers for the target gene was designed in GENEIOUS PRIME for the E4 esterase: Akond_E4_qPCR_F, CCTCGTCTTGGGTAACACTCA, and Akond_E4_qPCR_R, CATTTTCGCCGTTTGGTCCA. The aphid actin gene provided a reference gene with primers (Yang et al. 2023): actin_aphid_F1, GTGATGGTGTATCTCACACTGTC; and actin_aphid_R1, AGCAGTGGTGGTGAAACTG. Our qPCR reactions were performed according to Lee et al. (2012) with a Roche LightCycler 480. Briefly, 10-minute incubation at 95°C was used to heat activate the quick start IMMOLASE (Meridian, BIO-21047; Cincinnati, OH, USA), which was followed by 45 cycles of 95°C for 10 seconds, 54°C for 20 seconds, and 72°C for 15 seconds. The cycle ended with a high-resolution melt of 95°C for 1 minute (ramp rate = 4.8°C/second), 40°C for 1 minute (ramp rate = 2.5°C), and then an increase to 65°C (ramp rate = 4.8°C/second).

Fluorescence data was collected continuously from 65°C to 95°C (ramp rate = 0.02°C/second, 25 acquisitions per °C). Prior to our main experiment, we validated the amplification efficiency of primers using the Kellerberrin strain (Supplementary Figure S4).

A mean C_P_ (crossing-point) value for each aphid was calculated over two replicate qPCR reactions. We calculated the –ΔC_P_ as –(E4-like esterase C_P_ – actin C_P_) for each aphid. The 2^-ΔΔCP^ method (Livak and Schmittgen 2001) was used to estimate the relative expression (fold-change) between resistant and susceptible strains. We calculated the 2^-ΔΔCP^ score between each pair of resistant vs susceptible strains within each experiment. Within a strain comparison, we calculated 2^-ΔΔCP^ between all pairs of aphids from both strains. This allowed us to obtain a median value with 2.5% and 97.5% percentiles for the estimated 2^-ΔΔCP^.

#### Sequence variation

We characterized the distribution of segregating polymorphism in the exonic regions of our candidate E4-like esterase. For the eight core strains, we used SNPs that were not filtered for missing data and determined the protein coding effects through custom R code, combining the R packages BIOSTRINGS, DATA.TABLE (Dowle and Srinivasan 2019), TIDYVERSE (Wickham et al. 2019), SEQINR (Charif and Lobry 2007), and GENOMALICIOUS. We identified six non-synonymous SNPs that produced diagnostic genotypic differences between resistant and susceptible strains. Like our analysis of diagnostic mitochondrial SNPs (see above section ‘Clonal genetic background’), we developed a set of primers for the diagnostic non-synonymous E4-like esterase SNPs using GENEIOUS PRIME (see Supplementary Table S5). We then used these primers to genotype representative individuals from the field strains, using the same PCR protocol and analysis pipeline as derived from the diagnostic mitochondrial SNPs (see above section ‘Clonal genetic background’).

We performed protein sequence analysis to infer possible functional effects of the non- synonymous mutations. First, we aligned our *A. kondoi* E4-like esterase protein sequence against the *Myzus persicae* E4 esterase (UniProt Accession P35501) and the *Torpedo californica* acetylcholinesterase (UniProt Accession P04058) using GENEIOUS PRIME. We included the *T. californica* acetylcholinesterase because it is a model for type B carboxylesterases. Our alignments were used to understand whether the *A. kondoi* non- synonymous mutations occurred in or near the active site. Annotation of the active site was performed with the InterProScan server (Zdobnov and Apweiler 2001) as implemented on the EMBL-EBI server (https://www.ebi.ac.uk/interpro/search/sequence/). Second, we performed protein structural modelling of the *A. kondoi* E4-like esterase using the ALPHAFOLD 3 server (https://alphafoldserver.com/) (Abramson et al. 2024).

Quality of the predicted structure was assessed using ERRAT (Colovos and Yeates 1993) and PROCHECK (Laskowski et al. 1996) as implemented on the SAVES v6.1 server (https://saves.mbi.ucla.edu/).

### E4-like esterase transgenesis

Ectopic expression of our candidate *A. kondoi* E4-like esterase in *Drosophila melanogaster* (herein, ‘*Drosophila*’) was used to validate the role of this gene in insecticide resistance. Our analysis of sequence variation in the candidate E4-like esterase revealed non-synonymous differences between the susceptible and resistance *A. kondoi* strains (see Results). We were able to successfully obtain three strains of *D. melanogaster* carrying different E4-like esterase haplotypes, which we call H1, H2, and H3. H1 was identical to the susceptible reference genome sequence. H2 contained the mutations, Met150Leu and Pro586 Leu, which are heterozygotes in susceptible *A. kondoi* and homozygous in resistant *A. kondoi*. H3 contained three mutations, Ala28Val, Ala61Thr, and Ala527Thr, which are absent in the susceptible *A. kondoi*, and homozygous in the resistant *A. kondoi*.

The E4-like esterase was synthesised (Twist Bioscience) and cloned into the pUASTattB plasmid between the EcoRI and XbaI sites. (GenBank: EF362409.1). The construct was then transformed into the *Drosophila* germline, which carried an attP docking site on chromosome 2 (attP40) and the phiC31 integrase gene under the control of the vasa regulatory region on the X chromosome (y w M (eGFP, vas-int, dmRFP)ZH-2A: P [CaryP]attP40). Transgenic strains were balanced, and the integration of the transgenes confirmed using PCR and sequencing.

To conduct the bioassay, virgin females of the Act5C-GAL4 strain (“y[1] w[∗]; P(Act5C- GAL4-w)E1/CyO,””1;2”) (Bloomington Stock Center) were crossed with UAS-E4-like males to allow mating and oviposition. We exposed adult female flies to size concentrations of technical grade pirimicarb, chlorpyrifos and alpha-cypermethrin. The insecticide solutions were diluted in water and overlaid on 1.5% agar (1% sucrose, 0.5% acetic acid) in standard fly vials, then allowed to dry at room temperature overnight. Twenty adult females (2 to 7 days post-eclosion) were added to each vial and mortality assessed after 48 hours (alpha-cypermethrin) or 72 hours (pirimicarb and chlorpyrifos). Five replicates were conducted for each concentration. A *Drosophila* strain of the same genetic background as the two transgenic strains but lacking a transgene (progeny of crosses of the Act5C-GAL4 and UAS-no transgene strains) was included in all bioassays as a reference control. Control mortality was assessed using vials containing agar but lacking insecticide.

Our bioassays were performed as two experiments. We initially acquired the H2 mutant line before H1 and H3. In our first bioassay, Experiment 1, we contrasted the H2 mutant against the Control line. We were later able to acquire the H1 and H3 mutant lines.

Unfortunately, in the time between experiments, the H2 mutant line was lost. In our second bioassay, Experiment 2, we contrasted the H1 and H3 mutants against the

Control line. We acknowledge that running all three mutant lines in the same bioassay would have been ideal, but due to the circumstances, this was not possible. However, we note that Experiment 2 did contain lines carrying a susceptible version (H1) and a resistant version (H3) of the E4-like esterase, which provided a functional contrast of haplotypes carried in susceptible and resistant *A. kondoi*.

Dose-response data was analysed in R. For each insecticide and *Drosophila* strain combination, we fitted a quasibinomial model with mortality as a function of the log10- transformed insecticide concentration, using the GLM function from the STATS package. We derived χ^2^ goodness-of-fit test statistics and *p*-values (Brown 1978) using custom code. LD50-values were obtained with the DOSE.P function from the MASS package.

Resistance ratios for the mutant strain were calculated as the mutant LD50-value divided by the control LD50-value for each insecticide. Models were fit separately for each experiment.

### Target-site mutations

We used our combined transcriptome and whole-genome data to characterise allelic variation in target genes for organophosphate and carbamate insecticides (the acetylcholinesterase gene, *ace*) and pyrethroid insecticides (the *para*-like voltage-gated sodium channel, *para*). Our assembly of the *A. kondoi* genome identified three genes with *ace* annotations, with gene IDs, g11959, g46, and g15835, and lengths of 822, 706 and 664 amino acids, respectively. There were two genes with *para* annotations, with gene IDs, g11612 and g11614, and lengths of 1,160 and 958 amino acids, respectively. Sequence analysis was performed in R. To determine the relative codon positions of the target gene proteins, we aligned the *A. kondoi* sequences to their respective canonical reference species: *ace* sequences to *Torpedo californica* (X03439) and *para* sequences to *Musca domestica* (NP_001273814). Protein alignments were performed using the MSA v1.36.1 package (Bodenhofer et al. 2015).

We next determined whether non-synonymous mutations in our target genes might be associated with insecticide resistance. Here, we used SNPs that were not filtered for missing data and considered all possible alleles (not just biallelic loci) that occurred in the exonic regions of the annotated *ace* and *para* genes. Protein coding annotations were made with SNPEFF (Cingolani et al. 2012). These were then imported into R, where we extracted information about relative codon positions from our *A. kondoi* alignments to canonical reference proteins and matched these to our protein coding annotations using GENOMALICIOUS. We were most interested in any non-synonymous mutations in the primary *A. kondoi ace* and *para* gene, especially if they had been previously reported in the literature as conferring resistance.

We only found mutations in one of the *para* genes (see Results). To better contextualise these mutations, we analytically validated the functional roles of these annotated *para* genes. Aphids have evolved a heterodimeric *para* sodium channel, controlled by two genes, subunit 1 (domains I and II) and subunit 2 (domains III and IV) (Amey et al. 2015).

We therefore compared our *A. kondoi para* genes to two other aphids, the congeneric *A. pisum* and *M. persicae*. We used the *Myzus persicae* protein sequences of *para* subunits 1 and 2 (NCBI Protein Accessions CBI71141.1 and CBI71142.1, respectively) (Amey et al. 2015) to query the *Acythrosiphon pisum* genome (NCBI Genome Accession GCF_005508785.1) in NCBI’s BLASTP platform. We then recovered two respective top hits, with NCBI Protein Accession XP_029346126.1 (Gene ID 100158802) and XP_001949648.2 (Gene ID 100164620). In R, we aligned these *M. persicae* and *A. pisum* protein sequences with our *A. kondoi* sequences and that of the canonical *M. domestica* reference sequence, using MSA, visualizing the alignment with GENOMALICIOUS.

## Acknowledgements

We thank Alex Gill from the University of Melbourne for his advice and support on culturing *A. kondoi* in the lab. This work was supported by funding through the University of Melbourne’s ‘Early Carer Researcher Global Collaborations Award’ (JAT), and investment from the Grains Research and Development Corporation (UOM1906- 002RTX; UOM2404-006RT) and Hort Innovation (ST23002) through the ‘Australian Grain and Horticulture Pest Innovation Program’ (AAH and PAU) with additional support provided by the University of Melbourne and Cesar Australia. AgriFutures Australia provided additional financial support for sample collection (EC and PAU). The authors would like to acknowledge the use of the University of Exeter’s Advanced Research Computing facilities in carrying out this work. For the purpose of open access, the authors have applied a ‘Creative Commons Attribution (CC BY) licence to any Author Accepted Manuscript version arising from this submission.

## Contributions

### University of Melbourne

JAT, PAU, and AAH conceptualised the project. JAT developed the isofemale strains of *Acythosiphon kondoi* used in this study. JAT prepared all samples for transcriptomic and genomic sequencing. JAT co-led the nuclear genome assembly and differential expression analysis. JAT led the mitogenome assembly, analysis of clonal genetic backgrounds, experimental design and analysis of E4-like esterase expression and copy number validation, analysis of target-site mutations, analysis of diagnostic mitogenome and E4-like esterase SNPs, and analysis of dose-responses of transgenic *Drosophila*. CJB led the development of molecular protocols for validating E4- like esterase expression and copy number. RS led the molecular protocols for genotyping diagnostic mitochondrial and E4-like esterase SNPs and contributed to their analysis. Culturing and (or) molecular work associated with validating expression and copy number was contributed to by CJB, LAS, MS, KR, CJ, APSD, and QY. The original version of this manuscript was written by JAT, with contributions from AAH and PAU, and additional comments made by RS, CJB, LAS, MS, KR, CJ, APSD, and QY. AAH, JAT and PAU and acquired funding and performed project administration.

### University of Exeter

BJH co-led the genome assembly and differential expression analysis. SW and BT designed and executed the chemical bioassays of transgenic *Drosophila*. BT led the protein structural analysis. Contributions to original manuscript was made by CB, with additional comments made by BJH, BT, and SW. CB contributed funds for sequencing and hosted JAT as a visiting fellow at the University of Exeter.

### Cesar Australia

EC performed the original field collections of *A. kondoi*, identified the discriminating doses used to separate resistant from susceptible clones. MB contributed to molecular work associated with validating expression and copy number. Additional comments to the manuscript were made by EC and MB. EC and PAU acquired funds for this work.

## Data availability

The nuclear reference genome for *Acrythosiphon kondoi* that was generated in this study has been deposited into GenBank Genomes (JAUMJG000000000), and the partial mitochondrial genome has been submitted to GenBank Nucleotides (PV793158).

Genomic and transcriptomic reads associated with this work have been submitted as a GenBank BioProject PRJNA989388. The GenBank SRA accessions for sequencing reads, and Nucleotide accessions for Sanger sequences, can be found in Appendix 2. Custom gene annotations and counts of repetitive element classes that accompany our nuclear reference genome can be found in Appendix 3. Data and scripts associated with this work are archived through the University of Melbourne’s FigShare server (Thia 2025): doi.org/10.26188/29466431.

---

**Figure S1.**
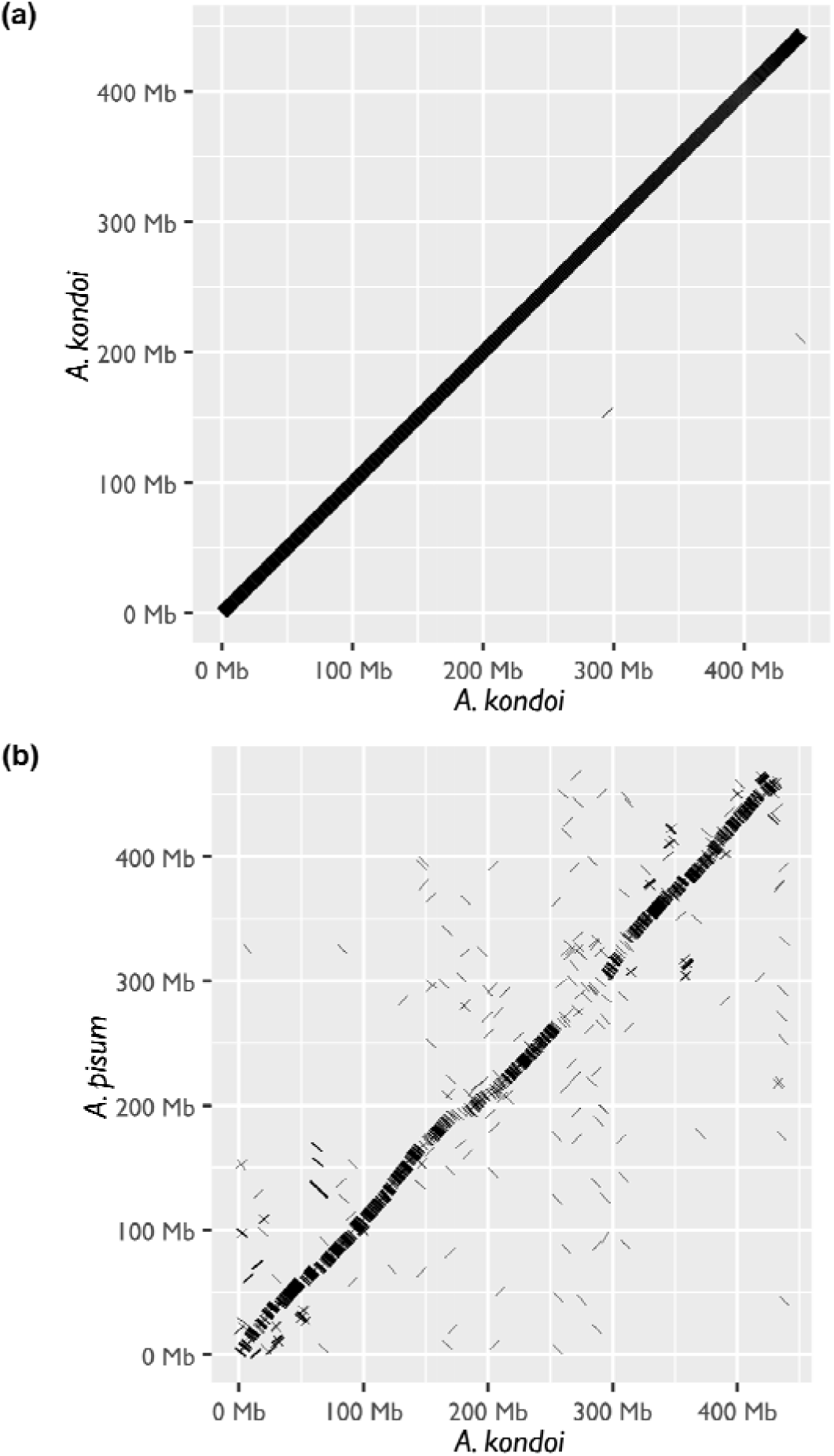
Dotplots of genomic alignments. The x-axis and y-axis indicate the concatenated genomic position (sequences lined up end-to-end). (a) Self-to-self alignment of our scaffold-level assembly of Acythorsiphon kondoi. The linear dotplot suggests that all sequences included in our final assembly are highly unique. (b) Alignment of A. kondoi pseudo- chromosomes to the A. pisum chromosome-level assembly (NCBI GCF_005508785.2). Acythosiphon kondoi pseudo-chromosomes were generated with RagTag in an initial alignment against the *A. pisum* genome. Derived pseudo-chromosomes were then mapped back to the *A. pisum* genome. (See also Figure S2)

**Figure S2.**
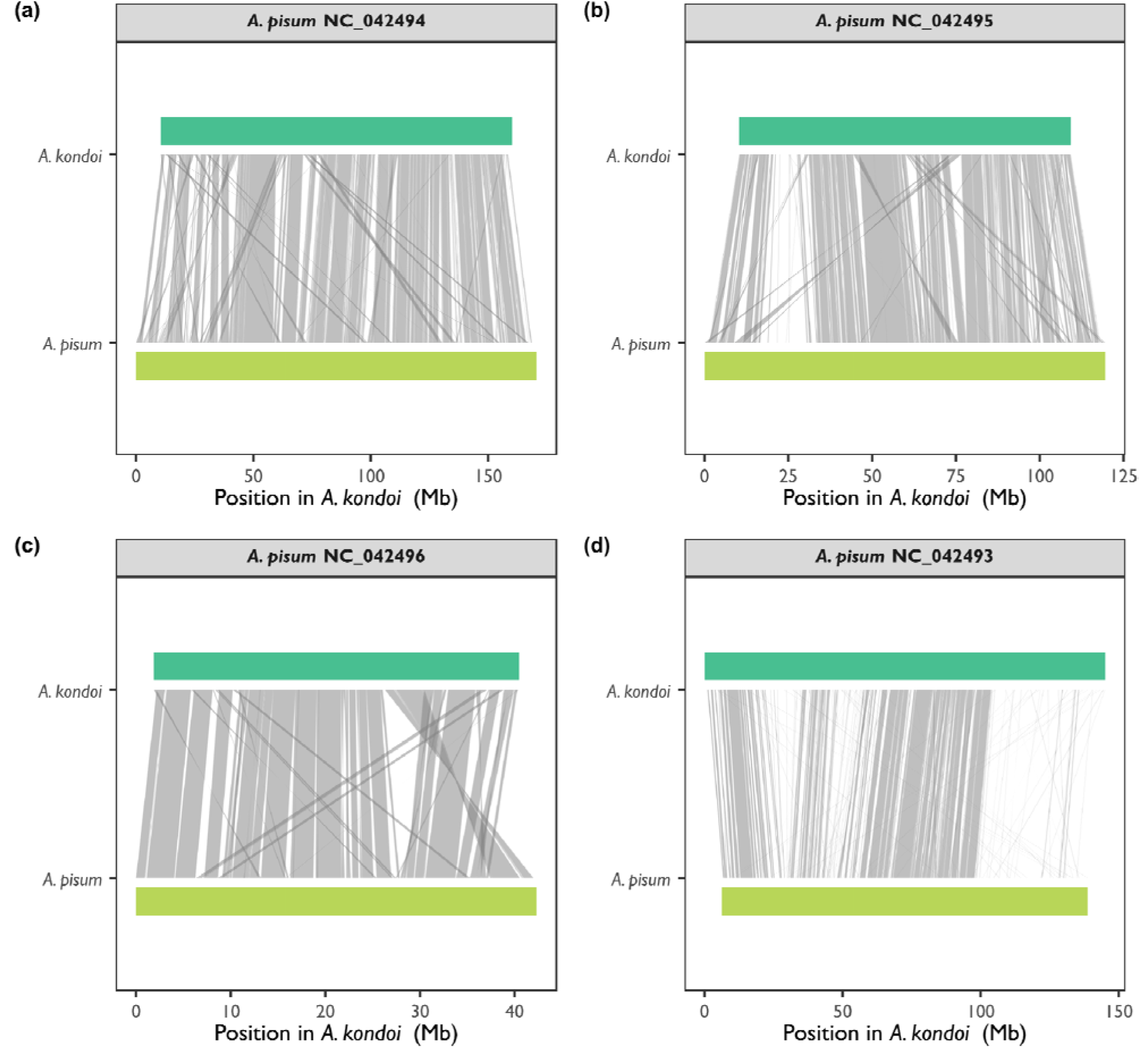
Realignment of *Acyrthoisphon kondoi* pseudo-chromosomes against the chromosome-level assembly of the *A. pisum* genome (NCBI GCF_005508785.2). The *x*-axis represents the position on the A. kondoi pseudo-chromosome. The *y*-axis delineates *A. kondoi* and *A. pisum* sequences. Grey bars illustrate alignments > 50,000 bp between the two sequences. Panels separate the three *A. pisum* chromosomes: (a) NC_042494, the A1 chromosome; (b) NC_042495, the A2 chromosome; (c) NC_042496, the A3 chromosome; and (d) NC_042493, the X chromosome. (See also Figure S1)

**Figure S3.**
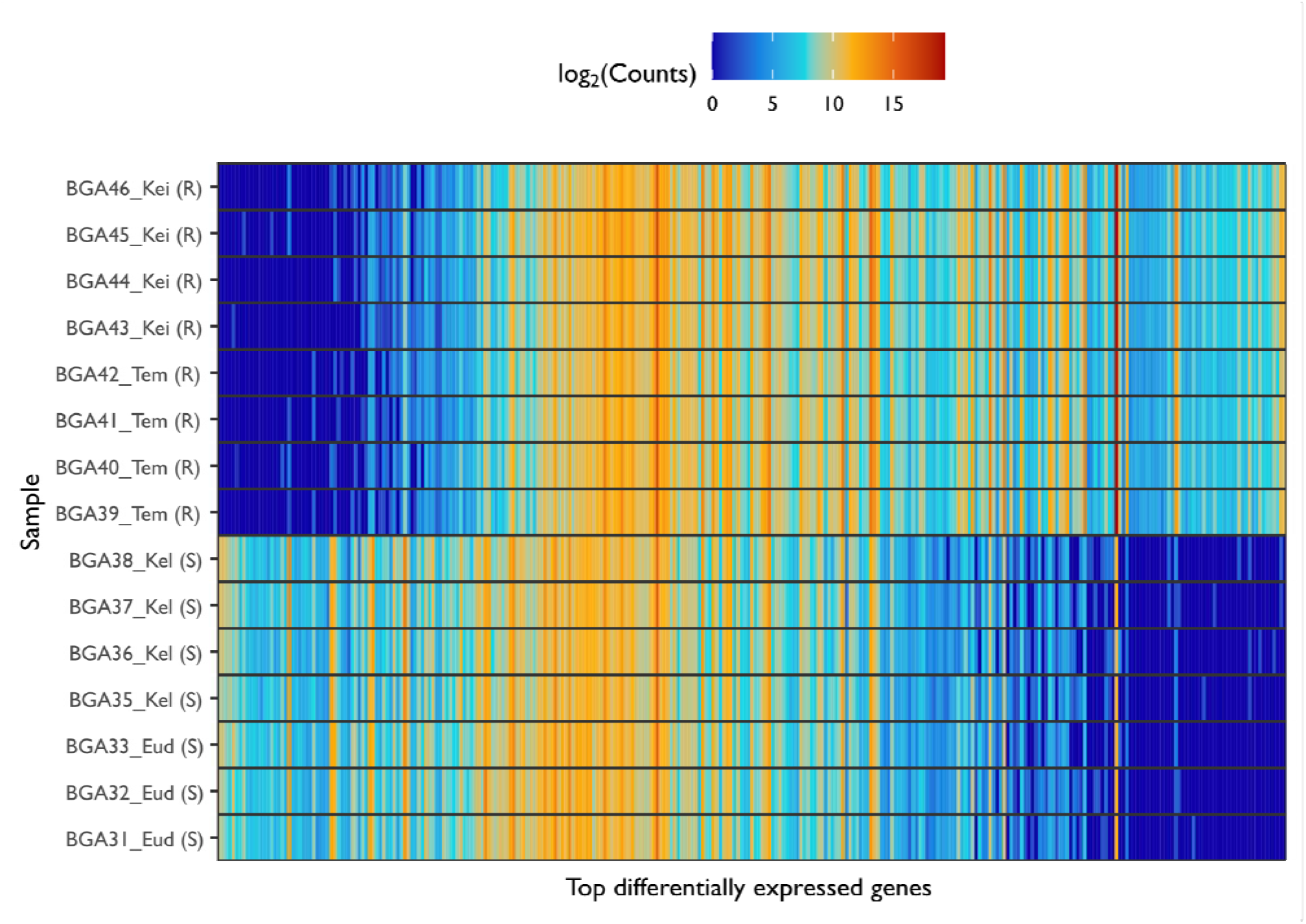
Read counts for the top differentially expressed genes (under- and over-expressed). Genes are on the x-axis, arranged from most under-expressed to most over-expressed in the resistant strains, relative to the susceptible strains. The y-axis represents different samples, labelled as ‘ID_Strain (Status)’: ‘ID’ is a unique sample identifier; ‘Strain’ is strains name; and ‘Status’ is one of ‘S’ for susceptible, or ‘R’ for resistant. Cell colours indicate the read counts (see legend).

**Figure S4.**
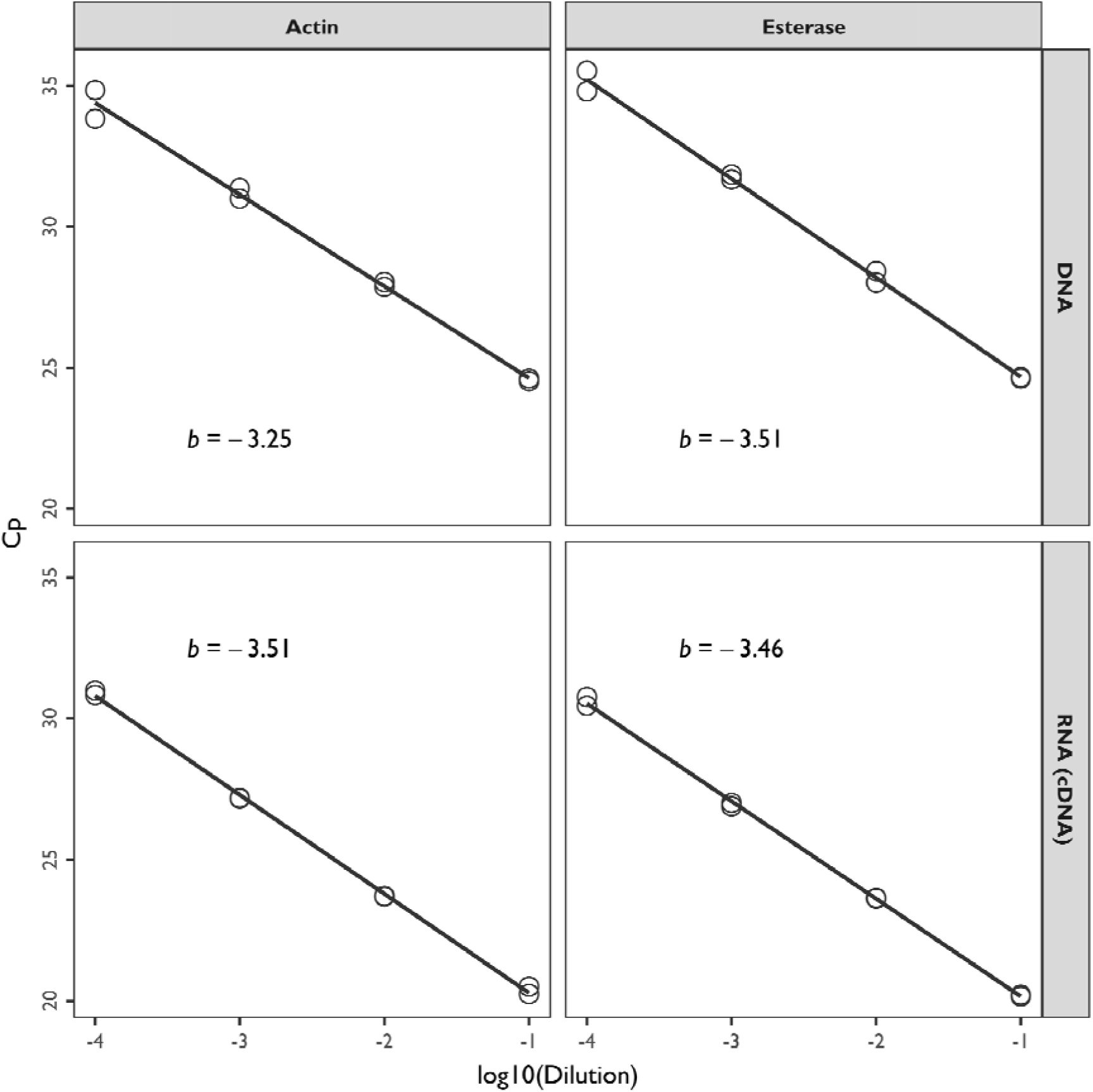
Primer efficiency test for estimating relative expression of the E4-like esterase. The x-axis represents the log10-transformed dilution factor of nucleic acid, whereas the y-axis represents the mean Cp value obtained in qPCR. Points represent different samples. Black lines indicate the line of best fit. The slope of the line, *b*, is also labelled on the plots: values between −3.3 and −3.6 are good. Panels separate results for the nucleic acid origin (in rows: DNA or RNA) and gene (in columns: actin or the esterase). These efficiency tests were performed on extractions prepared from the Kellerberrin strain.

**Figure S5.**
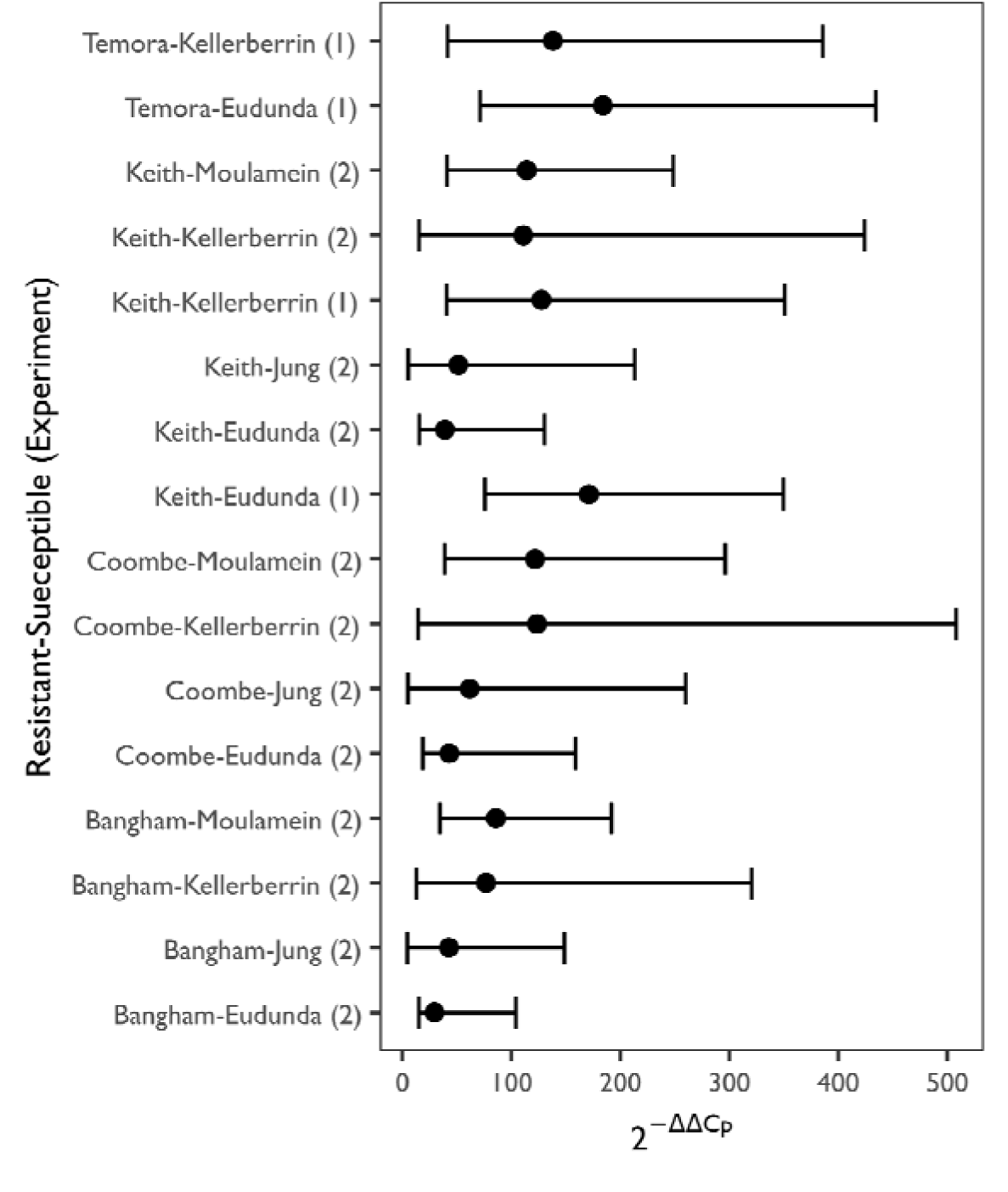
E4-like esterase fold-change expression differences between resistant vs. susceptible strains of *Acyrthosiphon kondoi*. The *x*-axis represents the measured fold-change difference, and the *y*-axis represents comparisons between different strains. Labels on the *y*-axis are presented as resistant vs susceptible with experimental number in parentheses. Points indicate median estimates of fold-change expression differences, with error bars indicating the 2.5% and 97.5% confidence intervals.

**Figure S6.**
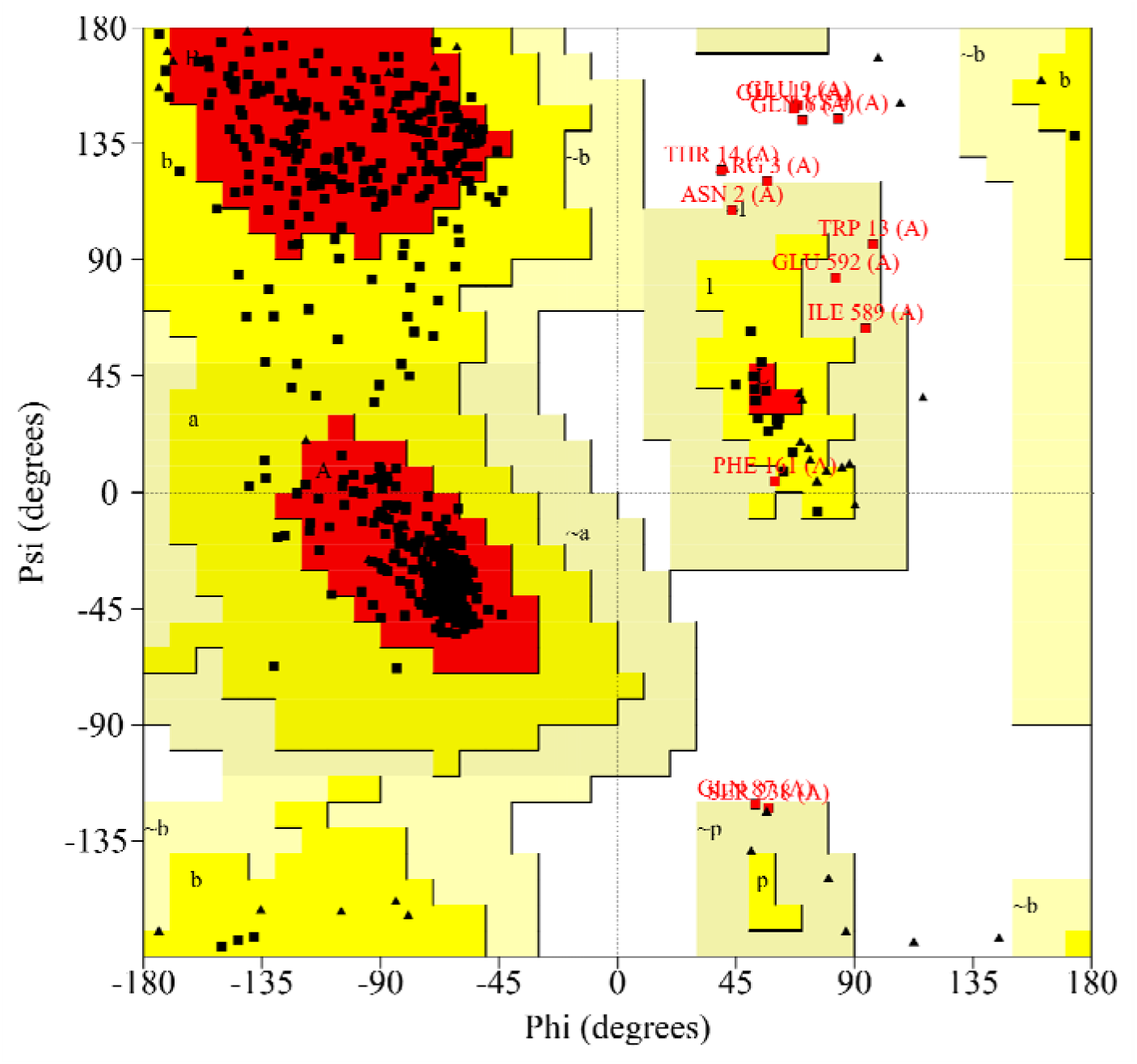
Ramachandran plot for our structural model of the *Acyrthosiphon kondoi* E4-like esterase. The *x-*axis represents the torsion angle around the N–C_α_ bond, and the y-axis represents the torsion angle around C_α_–C bonds Points represent amino acids, with red points indicating conformations in disallowed regions. Red shading indicates favourable conformations, yellow shading indicates allowed conformations, and white shading indicates disallowed conformations.

**Figure S7.**
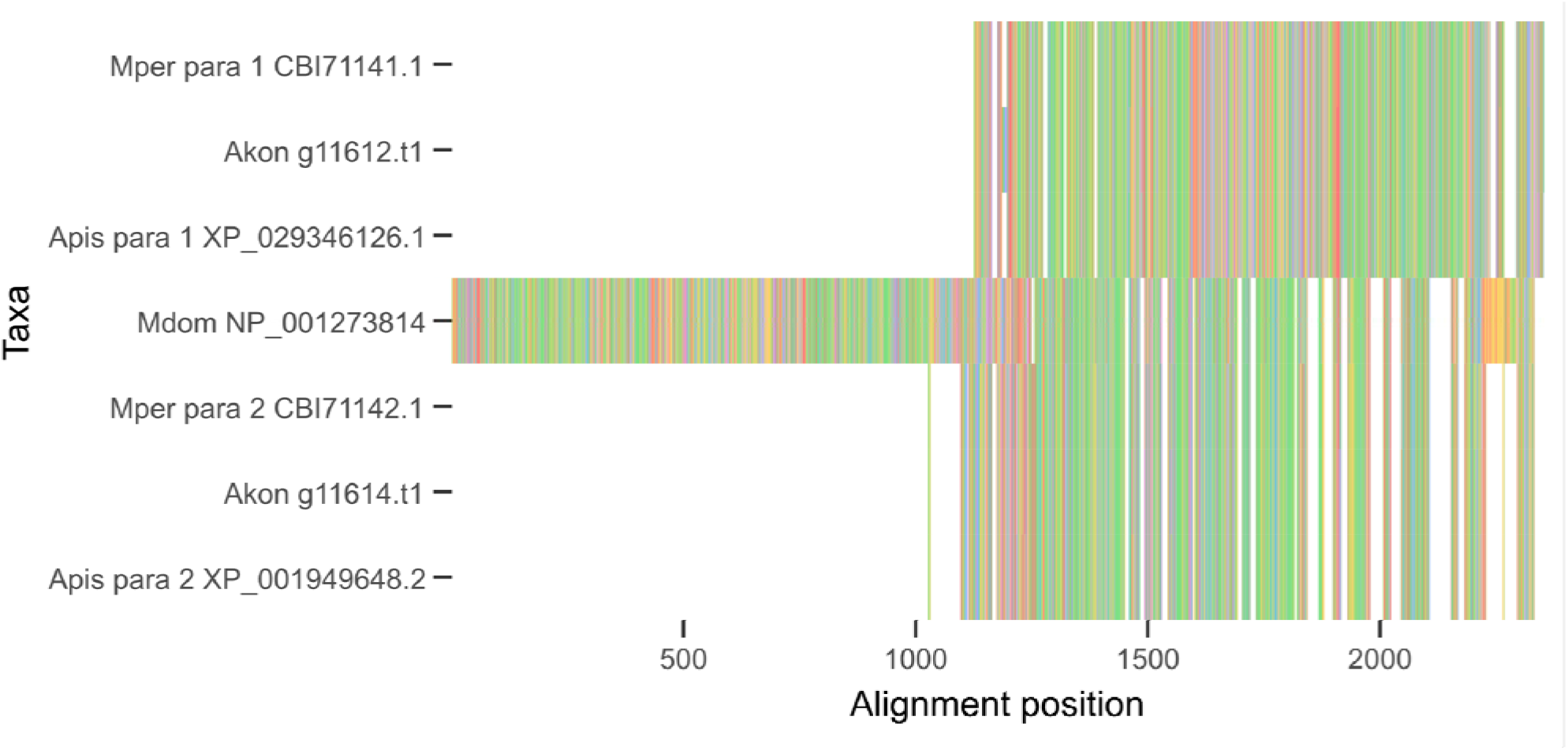
Alignment of *Acrythosiphon kondoi* (Akon) annotated *para* proteins to the canonical reference sequence, *Musca domestica* (Mdom), and to subunit 1 and 2 sequences found in the aphids *Myzus persicae* (Mper) and *A. pisum* (Apis). The alignment position is on the *x*-axis with species arranged on the *y*-axis. Colours represent the different amino acid classes.

**Figure S8.**
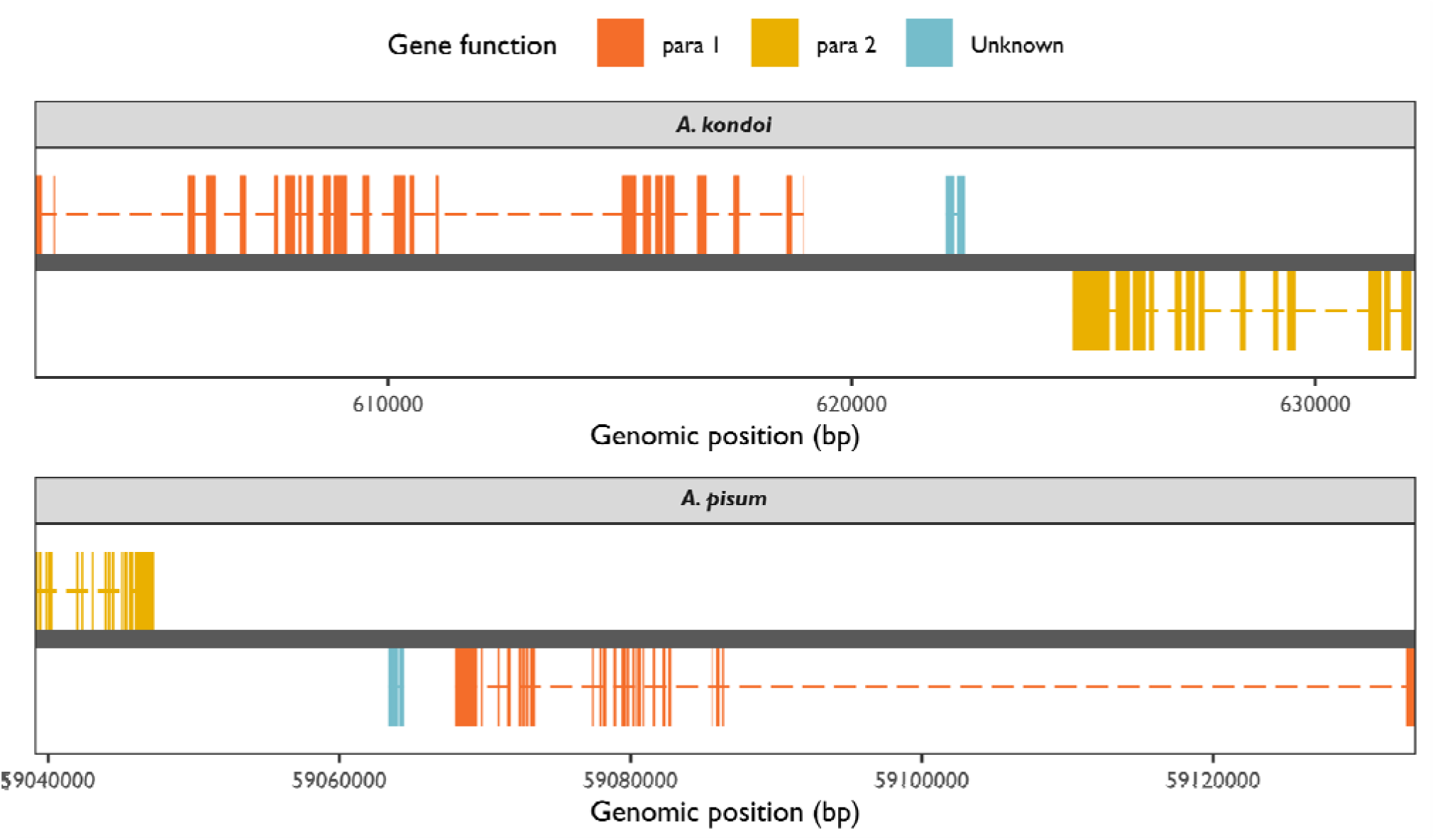
Genomic regions containing para genes in our reference assembly of *Acyrthosiphon kondoi* and in the reference assembly of *A. pisum* (NCBI Genome Accession GCF_005508785.1). The x-axis represents the genomic position in base pairs. The dark grey horizontal bar represents the genome. Coloured bars represent the exons of different genes, those on top represent genes on the positive strand, whereas those on the bottom represent genes on the negative strand. Dashed lines connect exons within the same gene. Exons are coloured by their gene function (see legend).

**Figure S9.**
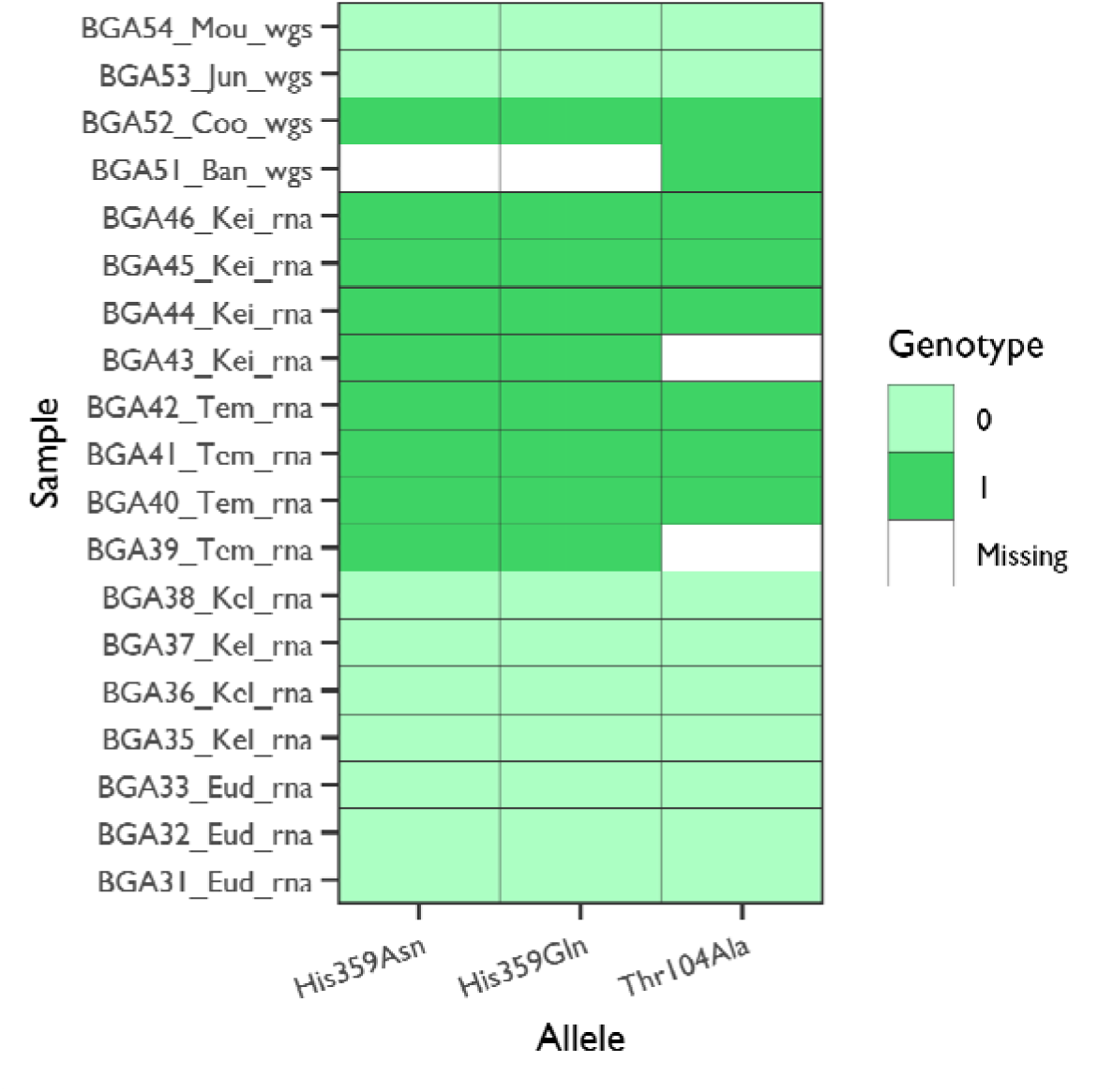
Non-synonymous mutations in the para g11614 gene annotated in the genome of *Acyrthosiphon kondoi*. Mutant alleles are arranged along the *x*-axis. Samples are arranged on the *y*-axis, labelled as ‘ID_Strain_Method’, where ‘ID’ is a unique sample identifier, ‘Strain’ is one of the eight clonal strains, and ‘Method’ is one of ‘rna’ for transcriptome sequencing or ‘wgs’ for whole-genome sequencing. Cell colours indicate the observed genotype (see legend). The susceptible strains include Eudunda (Eud), Kellerberrin (Kel), Jung (Jun) and Moulamein (Mou). The resistance strains include Bangham (Ban), Coombe (Coo), Keith (Kei) and Temora (Tem).

**Table S1.**
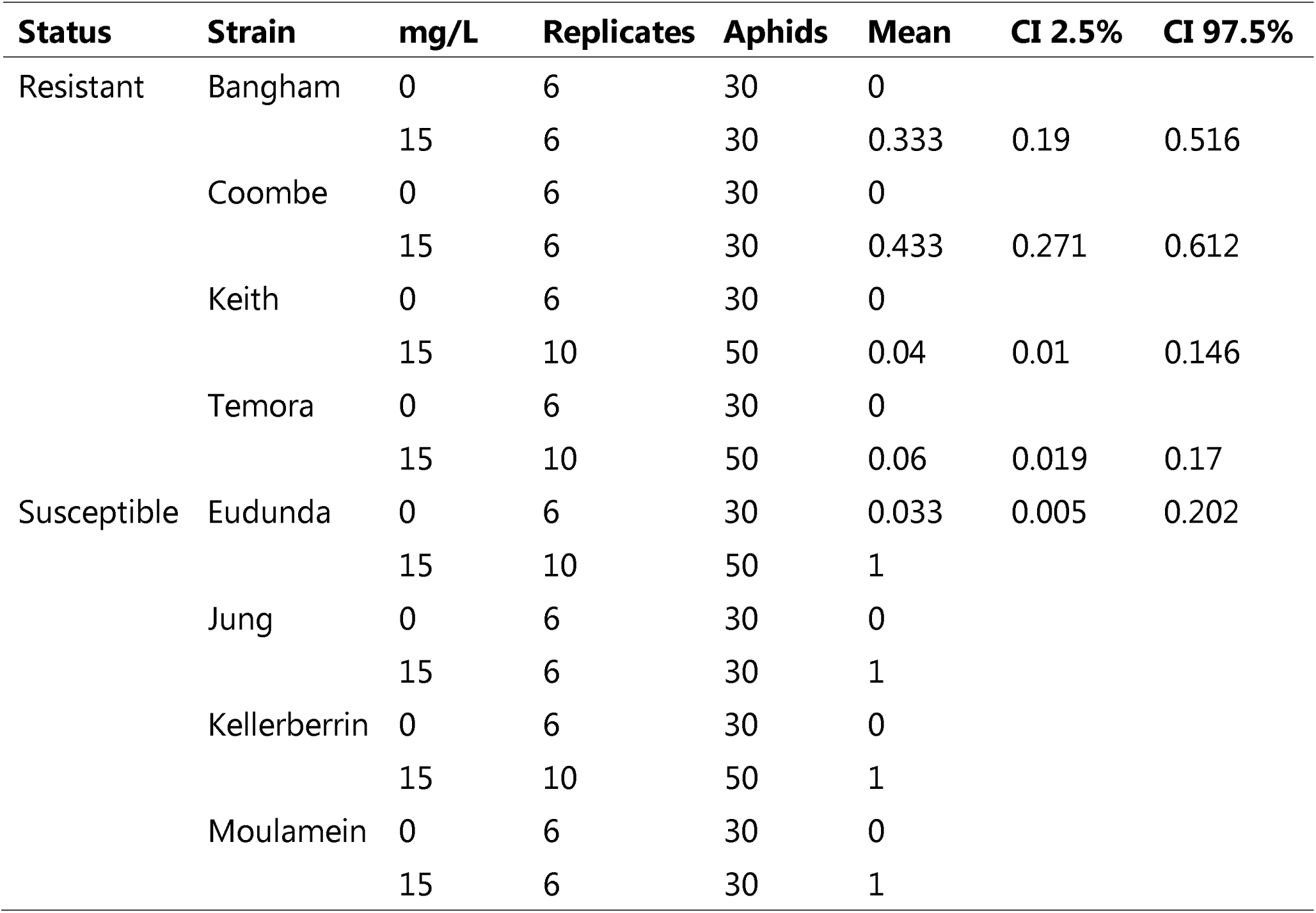
Mortality of the core *Acyrthosiphon kondoi* isofemale clonal strains to a water control and treatment of 15 mg a.i./L of chlorpyrifos in a leaf dip bioassay (72 hours exposure).

| Status | Strain | mg/L | Replicates | Aphids | Mean | CI 2.5% | CI 97.5% |
| --- | --- | --- | --- | --- | --- | --- | --- |
| Resistant | Bangham | 0 | 6 | 30 | 0 |  |  |
|  |  | 15 | 6 | 30 | 0.333 | 0.19 | 0.516 |
|  | Coombe | 0 | 6 | 30 | 0 |  |  |
|  |  | 15 | 6 | 30 | 0.433 | 0.271 | 0.612 |
|  | Keith | 0 | 6 | 30 | 0 |  |  |
|  |  | 15 | 10 | 50 | 0.04 | 0.01 | 0.146 |
|  | Temora | 0 | 6 | 30 | 0 |  |  |
|  |  | 15 | 10 | 50 | 0.06 | 0.019 | 0.17 |
| Susceptible | Eudunda | 0 | 6 | 30 | 0.033 | 0.005 | 0.202 |
|  |  | 15 | 10 | 50 | 1 |  |  |
|  | Jung | 0 | 6 | 30 | 0 |  |  |
|  |  | 15 | 6 | 30 | 1 |  |  |
|  | Kellerberrin | 0 | 6 | 30 | 0 |  |  |
|  |  | 15 | 10 | 50 | 1 |  |  |
|  | Moulamein | 0 | 6 | 30 | 0 |  |  |
|  |  | 15 | 6 | 30 | 1 |  |  |

**Table S2.** SNP heterozygosity for samples generated using transcriptome (‘rna’) and whole-genome (‘wgs’) sequencing at biallelic SNPs used to construct genetic distances.

| **Strain** | **Sample (R or S)** | **SNP heterozygosity** |
| --- | --- | --- |
| Eudunda | BGA31_Eud_rna (S) | 0.514 |
| Eudunda | BGA32_Eud_rna (S) | 0.515 |
| Eudunda | BGA33_Eud_rna (S) | 0.514 |
| Jung | BGA53_Jun_wgs (S) | 0.527 |
| Kellerberrin | BGA35_Kel_rna (S) | 0.494 |
| Kellerberrin | BGA36_Kel_rna (S) | 0.495 |
| Kellerberrin | BGA37_Kel_rna (S) | 0.495 |
| Kellerberrin | BGA38_Kel_rna (S) | 0.496 |
| Moulamein | BGA54_Mou_wgs (S) | 0.527 |
| Bangham | BGA51_Ban_wgs (R) | 0.588 |
| Coombe | BGA52_Coo_wgs (R) | 0.589 |
| Keith | BGA43_Kei_rna (R) | 0.572 |
| Keith | BGA44_Kei_rna (R) | 0.572 |
| Keith | BGA45_Kei_rna (R) | 0.574 |
| Keith | BGA46_Kei_rna (R) | 0.574 |
| Temora | BGA39_Tem_rna (R) | 0.573 |
| Temora | BGA40_Tem_rna (R) | 0.574 |
| Temora | BGA41_Tem_rna (R) | 0.573 |
| Temora | BGA42_Tem_rna (R) | 0.573 |

**Table S3.** Fold-change difference in the expression of the candidate E4-like esterase between resistant and susceptible strains estimated from qPCR assays with the 2^-DDCP^ method. Median values with 2.5% and 97.5% percentiles for the estimates are reported.

| Experiment | Resistant | Susceptible | Median | 2.5% | 97.5% |
| --- | --- | --- | --- | --- | --- |
| 1 | Keith | Eudunda | 171 | 76 | 350 |
|  | Keith | Kellerberrin | 128 | 41 | 351 |
|  | Temora | Eudunda | 184 | 71 | 434 |
|  | Temora | Kellerberrin | 138 | 42 | 386 |
| 2 | Bangham | Eudunda | 29 | 15 | 104 |
|  | Bangham | Jung | 43 | 5 | 149 |
|  | Bangham | Kellerberrin | 77 | 13 | 321 |
|  | Bangham | Moulamein | 86 | 34 | 192 |
|  | Coombe | Eudunda | 43 | 19 | 159 |
|  | Coombe | Jung | 62 | 5 | 260 |
|  | Coombe | Kellerberrin | 124 | 15 | 508 |
|  | Coombe | Moulamein | 122 | 39 | 296 |
|  | Keith | Eudunda | 39 | 16 | 130 |
|  | Keith | Jung | 52 | 5 | 213 |
|  | Keith | Kellerberrin | 111 | 15 | 424 |
|  | Keith | Moulamein | 114 | 41 | 249 |

**Table S4.** Results from insecticide bioassays on transgenic *Drosophila melanogaster* carrying our candidate E4-like esterase from *Acyrthosiphon kondoi*. Reported are the intercepts (*b*_0_), slopes (*b*_1_), *χ*^2^ goodness-of-fit statistics and associated *P*-values, LD_50_ values (mg a.i./L) with 95% confidence intervals, and resistance ratios (RR) of mutant strains. (See also Figure 4)

| Experiment | Strain | Insecticide | $b_0$ | $b_1$ | $\chi^2$ (df) | $P(\chi^2)$ | LD50 values (95% CIs) | RR |
| --- | --- | --- | --- | --- | --- | --- | --- | --- |
| 1 | Mut-H2 | Pirimicarb | -10.61 | 4.05 | 0.60 (3) | 0.896 | 416.7 (369.71,469.66) | 3.19 |
|  |  | Chlorpyrifos | 1.79 | 4.39 | 0.41 (3) | 0.939 | 0.39 (0.36,0.43) | 4.88 |
|  |  | Alpha-cypermethrin | -2.09 | 3.76 | 0.65 (3) | 0.884 | 3.6 (3.16,4.11) | 4.29 |
|  | Control | Pirimicarb | -8.64 | 4.09 | 0.40 (3) | 0.94 | 130.5 (118.01,144.31) | – |
|  |  | Chlorpyrifos | 4.06 | 3.67 | 0.51 (3) | 0.916 | 0.08 (0.07,0.09) | – |
|  |  | Alpha-cypermethrin | 0.32 | 4.15 | 0.40 (3) | 0.94 | 0.84 (0.74,0.94) | – |
| 2 | Mut-H1 | Pirimicarb | -8.68 | 3.09 | 0.06 (3) | 0.997 | 637.96 (578.62,703.39) | 2.12 |
|  |  | Chlorpyrifos | 0.54 | 2.99 | 0.06 (3) | 0.996 | 0.66 (0.6,0.71) | 4.40 |
|  |  | Alpha-cypermethrin | -3.51 | 5.60 | 0.30 (3) | 0.961 | 4.24 (3.94,4.58) | 2.81 |
|  | Mut-H3 | Pirimicarb | -9.70 | 3.54 | 0.05 (3) | 0.997 | 545.54 (497.4,598.34) | 1.81 |
|  |  | Chlorpyrifos | 1.64 | 4.49 | 0.10 (3) | 0.991 | 0.43 (0.4,0.47) | 2.87 |
|  |  | Alpha-cypermethrin | -2.60 | 4.60 | 0.66 (3) | 0.883 | 3.67 (3.29,4.1) | 2.43 |
|  | Control | Pirimicarb | -11.97 | 4.83 | 0.53 (3) | 0.913 | 301.35 (270.88,335.26) | – |
|  |  | Chlorpyrifos | 6.12 | 7.34 | 0.23 (3) | 0.972 | 0.15 (0.14,0.15) | – |
|  |  | Alpha-cypermethrin | -1.21 | 6.78 | 0.09 (3) | 0.993 | 1.51 (1.42,1.61) | – |

**Table S5.**
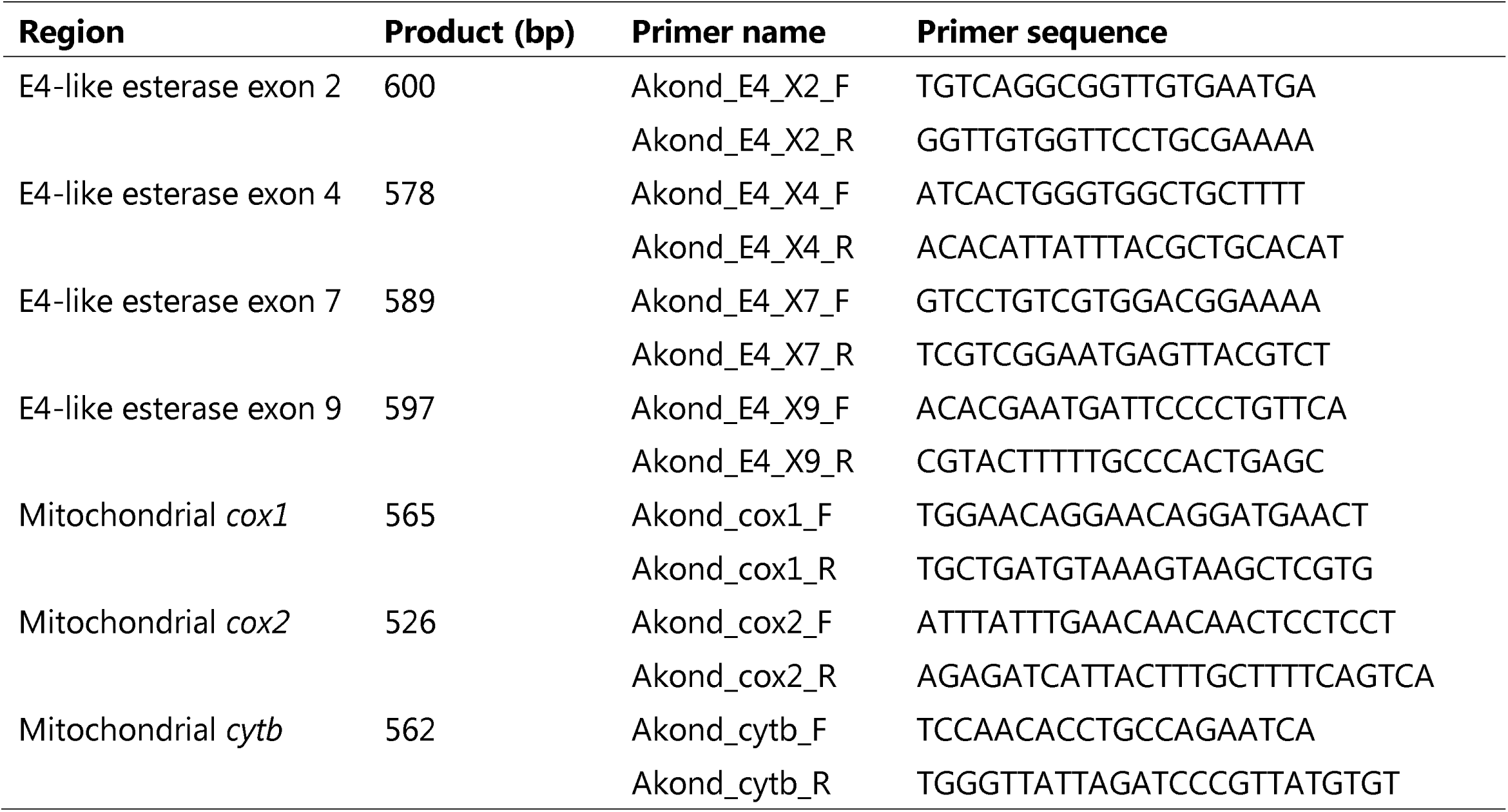
Primers for PCR and Macrogen sequencing of focal regions in the E4-like esterase and mitochondrial genes.

